# Molecular Basis of Antibiotic Self-Resistance in a Bee Larvae Pathogen

**DOI:** 10.1101/2021.11.23.469742

**Authors:** Tam Dang, Bernhard Loll, Sebastian Müller, Ranko Skobalj, Julia Ebeling, Timur Bulatov, Sebastian Gensel, Josefine Göbel, Markus C. Wahl, Elke Genersch, Andi Mainz, Roderich D. Süssmuth

## Abstract

*Paenibacillus larvae*, the causative agent of the devastating honey-bee disease American Foulbrood, produces the cationic polyketide-peptide hybrid paenilamicin that displays high antibacterial and antifungal activity. Its biosynthetic gene cluster contains a gene coding for the *N*-acetyltransferase PamZ. We show that PamZ acts as self-resistance factor in *P. larvae* by deactivation of paenilamicin. Using tandem MS, NMR spectroscopy and synthetic diastereomers, we identified the N-terminal amino group of the agmatinamic acid as the *N*-acetylation site. These findings highlight the pharmacophore region of paenilamicin, which we very recently identified as a new ribosome inhibitor. Here, we further elucidated the crystal structure of PamZ:acetyl-CoA complex at 1.34 Å resolution. An unusual tandem-domain architecture provides a well-defined substrate-binding groove decorated with negatively-charged residues to specifically attract the cationic paenilamicin. Our results will help to understand the mode of action of paenilamicin and its role in pathogenicity of *P. larvae* to fight American Foulbrood.

## Introduction

Pollination of wild and cultivated flowering plants is an indispensable ecosystem service, which is mainly provided by pollinating insects. Among the insect pollinators, managed honey bee colonies play a particularly important role in agriculture, where they are widely used as commercial pollinators and contribute to 35% of the production volume of global food crops^1^. In order to secure human food supply, it is therefore important to ensure the health of honey bees, which is continuously threatened by various viral, bacterial and fungal pathogens as well as metazoan parasites^2^.

The Gram-positive, facultative anaerobic, spore-forming bacterium, *Paenibacillus larvae* (*P. larvae*), is the causative agent of the epizootic American Foulbrood (AFB) of honey bees^3^. AFB is a fatal intestinal infection of the honey bee brood initiated in first instar larvae by ingestion of spore-contaminated food. The distribution of the spores, the infectious form of *P. larvae*, within a colony and between colonies, also within apiary and between apiaries^4^, consequently leads to honey bee colony losses. *P. larvae* comprises the four well-described genotypes ERIC I to ERIC IV^3^ which differ in virulence on the larval^5^ and colony level^6^ as well as in pathogenesis strategies employed to kill the host^7^. The existence of another ERIC genotype, ERIC V, has recently been proposed^8^. From contemporary outbreaks of AFB all over the world, only *P. larvae* ERIC I and ERIC II can be isolated^9^ suggesting that the hypervirulent genotypes ERIC III to ERIC V did not become established in the honey bee population.

In our quest to find sustainable control measures against this most serious bacterial disease of honey bees, we started to unravel AFB pathogenesis by analyzing the interaction between *P. larvae* and honey bee larvae on a molecular level. We identified several virulence factors of *P. larvae* ERIC I and ERIC II and showed that two AB toxins^10,11^, a chitin-degrading enzyme^12,13^ and also an S-layer protein^14,15^ have a pivotal role in the virulence of this pathogen and that *P. larvae* also produces various secondary metabolites^16^. Bacterial secondary metabolites, with polyketides and (non-)ribosomal peptides as important representatives, provide highly valuable lead structures, among them antibiotics with novel modes of actions for drug development to fight various infectious diseases^17,18^. Secondary metabolites can also act as virulence(-like) factors, functioning as signal molecules in gene regulation of defense or growth mechanisms^19–21^. The search for secondary metabolites produced by *P. larvae* led to the structural elucidation of paenilamicin that shows cytotoxic, antibacterial and antifungal activities^22,23^. It is currently assumed that paenilamicin is produced as a defense molecule against microbial competitors, since only *P. larvae* can be isolated as a pure culture from AFB-diseased larval cadavars, while other microbial competitors are absent in the degradation process of the infected larvae^24^.

Paenilamicin is a linear, cationic aminopolyol peptide antibiotic and is synthesized via an unusual nonribosomal peptide synthetase-polyketide synthase (NRPS-PKS) hybrid assembly line that exhibits several fascinating biosynthetic features. It contains unusual structural motifs such as galantinamic acid (Glm), agmatinamic acid (Aga), *N*-methyldiaminopropionic acid (mDap), galantinic acid (Gla) and a 4,3-spermidine (Spd) at the C-terminus (**Figure 1**). *P. larvae* produces a mixture of paenilamicin variants A1, A2, B1 and B2. They only differ in two positions of the paenilamicin backbone: at the N-terminus and in the center between mDap1 and Gla. Either a lysine (series A) or an arginine (series B) is activated by the adenylation domain of NRPS1 (**Figure 1**). The amino acid residue between mDap1 and Gla is a lysine (series 1) or an ornithine (series 2) assigned to be incorporated by NRPS4 (*pamD*), respectively (**Figure 1**).

**Figure 1.**
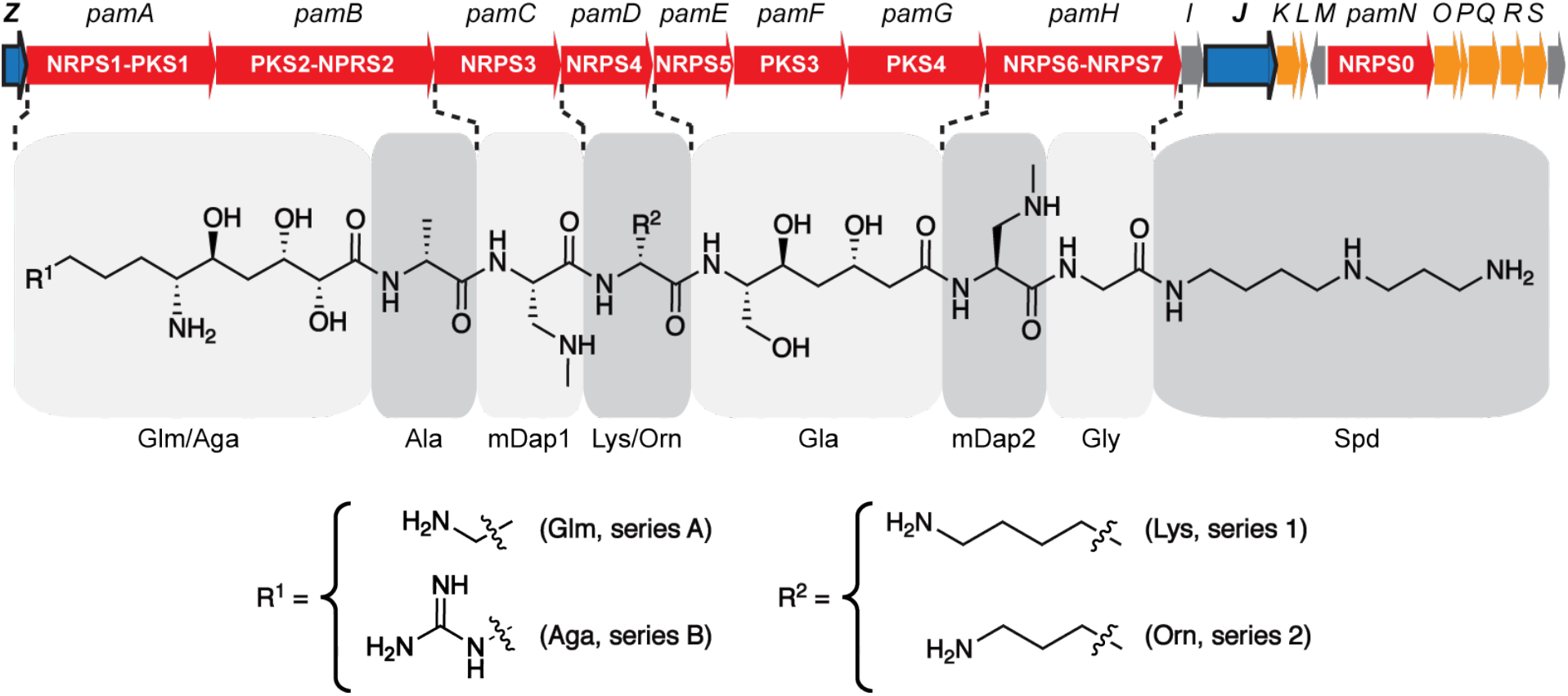
Biosynthetic gene cluster and structure of paenilamicin variants. The *pam* gene cluster^22^ contains core biosynthetic (red), auxiliary biosynthetic (orange), resistance (*pamZ* and *pamJ*; blue) and other (grey) genes and expresses the NRPS-PKS hybrid biosynthetic machinery for the production of paenilamicin A1 (Glm, Lys), A2 (Glm, Orn), B1 (Aga, Lys) and B2 (Aga, Orn). Abbreviations are listed as follows: galantinamic acid (Glm), agmatinamic acid (Aga), lysine (Lys), ornithine (Orn), alanine (Ala), *N*-methyldiaminopropionic acid (mDap), galantinic acid (Gla), glycine (Gly), 4,3-spermidine (Spd).

The *pam* gene cluster harbors a gene encoding the putative acetyl-CoA-dependent *N*-acetyltransferase, PamZ, which belongs to the Gcn5-related *N*-acetyltransferase (GNAT) superfamily^25,26^. One prominent member of this superfamily is the bacterial aminoglycoside *N*-acetyltransferase (AAC) that plays an important role in antibiotic resistances, particularly in clinical and environmental settings^27^. Aminoglycoside antibiotics have been widely used in the treatment of bacterial infections but they rapidly lose activity against multi-resistant bacteria due to adaptation and the development of resistance. By contrast, self-resistance is an innate, non-adaptation-based mechanism for the protection against self-produced antimicrobial agents. Since self-produced antimicrobial agents could also harm the bacterial host, self-resistance is critical for survival and territorial competition.

Our results demonstrate the deactivation of paenilamicins by the regio-and stereoselective self-resistance protein PamZ including its high-resolution crystal structure that shows how its tandem-domain arrangement organizes substrate binding. Together with a parallel study^28^, in which we report on the total synthesis and the biological evaluation of paenilamicin, we have here unambiguously identified the N-terminal building block of paenilamicins as an essential switch for target binding, biological activity and self-resistance.

## Results

### Regio- and stereoselective *N*-acetylation of paenilamicin by PamZ

To confirm our hypothesis that PamZ (NCBI accession no.: WP_023484187) is an acetyl-CoA-dependent *N*-acetyltransferase that targets paenilamicins, we monitored PamZ-mediated antibacterial effects *in vitro* by agar diffusion assays against *Bacillus megaterium* (*B. megaterium*) as indicator strain as well as by mass spectrometry (MS) and nuclear magnetic resonance (NMR) spectroscopy. To this end, the *pamZ* gene was amplified from the wild type (WT) *P. larvae* ERIC II strain, inserted into the commercial pET28a(+) vector and transformed into *E. coli* BL21-Gold(DE3) for heterologous expression. PamZ was then purified (**Figure S1**) and used for the assays including four native paenilamicin variants as substrates and acetyl-CoA as co-substrate. The paenilamicin variants were purified from *P. larvae* ERIC I and ERIC II, which preferably produce the paenilamicin mixtures A2/B2 and A1/B1, respectively (**Figure 1, Figure S2**). In addition, we also tested synthetic paenilamicin B2 (PamB2_3)^28^.

The agar diffusion assays clearly showed that paenilamicins incubated with PamZ and acetyl-CoA were not able to inhibit the growth of *B. megaterium*, whereas antibacterial activity was observed in the absence of acetyl-CoA and/or PamZ (**Figure 2**). This loss of biological activity correlated with the conversion of paenilamicins to the corresponding *N*-acetylpaenilamicins as observed by HPLC-ESI MS. ESI mass spectra revealed that the mass-to-charge ratios of natural and synthetic paenilamicins exhibited a characteristic mass shift of 42 Da indicative of the addition of an acetyl group (**Figure S3-S7**).

**Figure 2.**
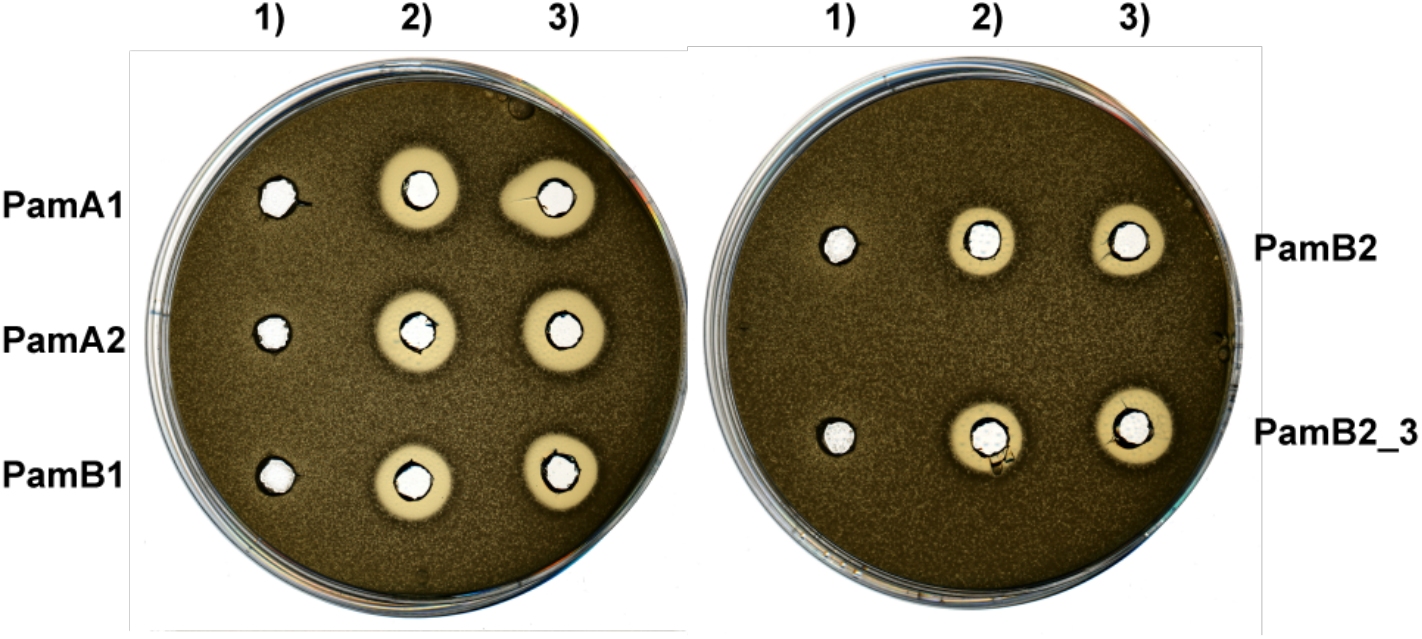
Deactivation of paenilamicins through PamZ-mediated *N*-acetylation tested by agar diffusion assay against *B. megaterium* as indicator strain. Paenilamicin variants (PamA1, A2, B1, B2) isolated from *P. larvae* and synthetic paenilamicin B2 (PamB2_3) were incubated *in vitro* with both acetyl-CoA and PamZ (1), acetyl-CoA only (2) or PamZ only (3). Samples 2 and 3 are negative controls and indicate the lack of bacterial growth.

Paenilamicin contains several primary and secondary amino groups that are potential candidates for *N*-acetylation. To determine the site of acetylation, we monitored PamZ-mediated effects in fingerprint tandem MS and NMR spectra of paenilamicin before and after treatment with PamZ/acetyl-CoA. Besides the mass shift of 42 Da for the acetylation, characteristic MS^2^ fragmentation patterns originated from the difference between Glm and Aga residues in series A and B (+28 Da) as well as the difference between Lys and Orn residues in series 1 and 2 (+14 Da). MS^2^ fragmentation mainly resulted in fragment ions b_4_, y_4_ and y_6_ of each paenilamicin and *N*-acetylpaenilamicin variant acquired by collision-induced dissociation (**Table S1**). Fragment ion b_4_ varied depending on the paenilamicin series showing mass shifts of 14 Da and 28 Da. Importantly, we observed a mass shift of 42 Da only for fragment ion b_4_, indicating acetylation in the N-terminal half of paenilamicin. By contrast, the fragment ions y_4_ and y_6_ did not exhibit any mass shifts of 42 Da between paenilamicins and *N*-acetylpaenilamicins. Thus, we excluded acetylation in the C-terminal half of paenilamicin (**Figure S8-S18**). In addition, we detected and isolated small amounts of *N*-acetylpaenilamicin A1, B1 and B2 from supernatants of *P. larvae* ERIC I and ERIC II (**Figure S19**), and compared them with our products formed *in vitro*. The MS^2^ fragmentation analysis confirmed that the mono-acetylation in the N-terminal half of paenilamicin also occurred *in vivo* (**Figure S20-S22**). The MS^2^ experiments did not reveal whether the N-terminal amino group of Aga-6 or its side chain (amino/guanidino group) was acetylated.

To ultimately identify the functional group that is modified by PamZ, we acquired ^1^H-^13^C hetero-nuclear single-quantum coherence (HSQC) NMR spectra of paenilamicin B2 before and after incubation with PamZ/acetyl-CoA. Although both spectra were mostly superimposable, severe chemical shift perturbations (CSPs) were observed for a minor fraction of cross-peaks (**Figure 3a**). Mapping CSPs onto the structure of paenilamicin B2 revealed a well-defined region comprising the N-terminal half, with the strongest effect being located at Aga-6 (**Figure 3b, Table S2**). *N*-acetylpaenilamicin B2 also showed an additional cross-peak compared to paenilamicin B2, which we tentatively assigned to the methyl moiety of the newly attached acetyl group (**Figure 3a**). Our data unequivocally demonstrated that PamZ mono-*N*-acetylates the N-terminal amino group at Aga-6 position of paenilamicin and thereby abolishes its antibacterial activity. Ultimately, this result is further supported by two synthetic diastereomers of paenilamicin B2 with L-instead of the native D-configuration at Aga-6 (PamB2_1 and PamB2_2), that were both antibacterially less active^28^ and that were not modified by PamZ (**Figure 4, Figure S23**).

**Figure 3.**
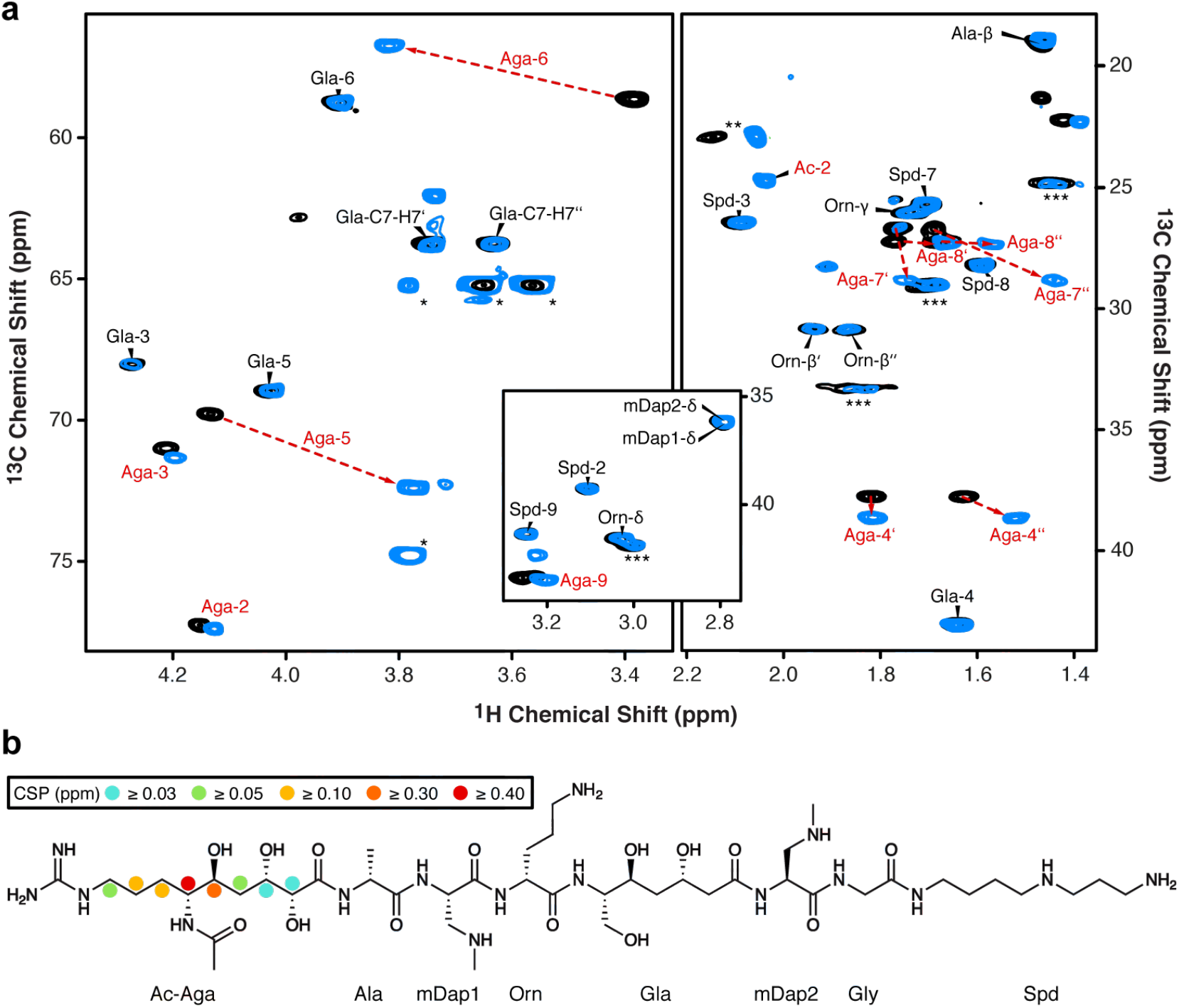
Identification of the *N*-acetylation site through 2D NMR spectroscopy. **a** Overlay of relevant ^1^H-^13^C HSQC sections of paenilamicin B2 (black) and *N*-acetylpaenilamicin B2 (blue). Strongly-perturbed cross-peaks are highlighted with red labels. Known impurities are labeled with one, two and three asterisks arising from glycerol, acetic acid and residual purification traces of paenilamicin B1, respectively. **b** Significant chemical shift perturbations (CSPs) are indicated as circles (see legend for color code) in the chemical structure of *N*-acetylpaenilamicin B2.

**Figure 4.**
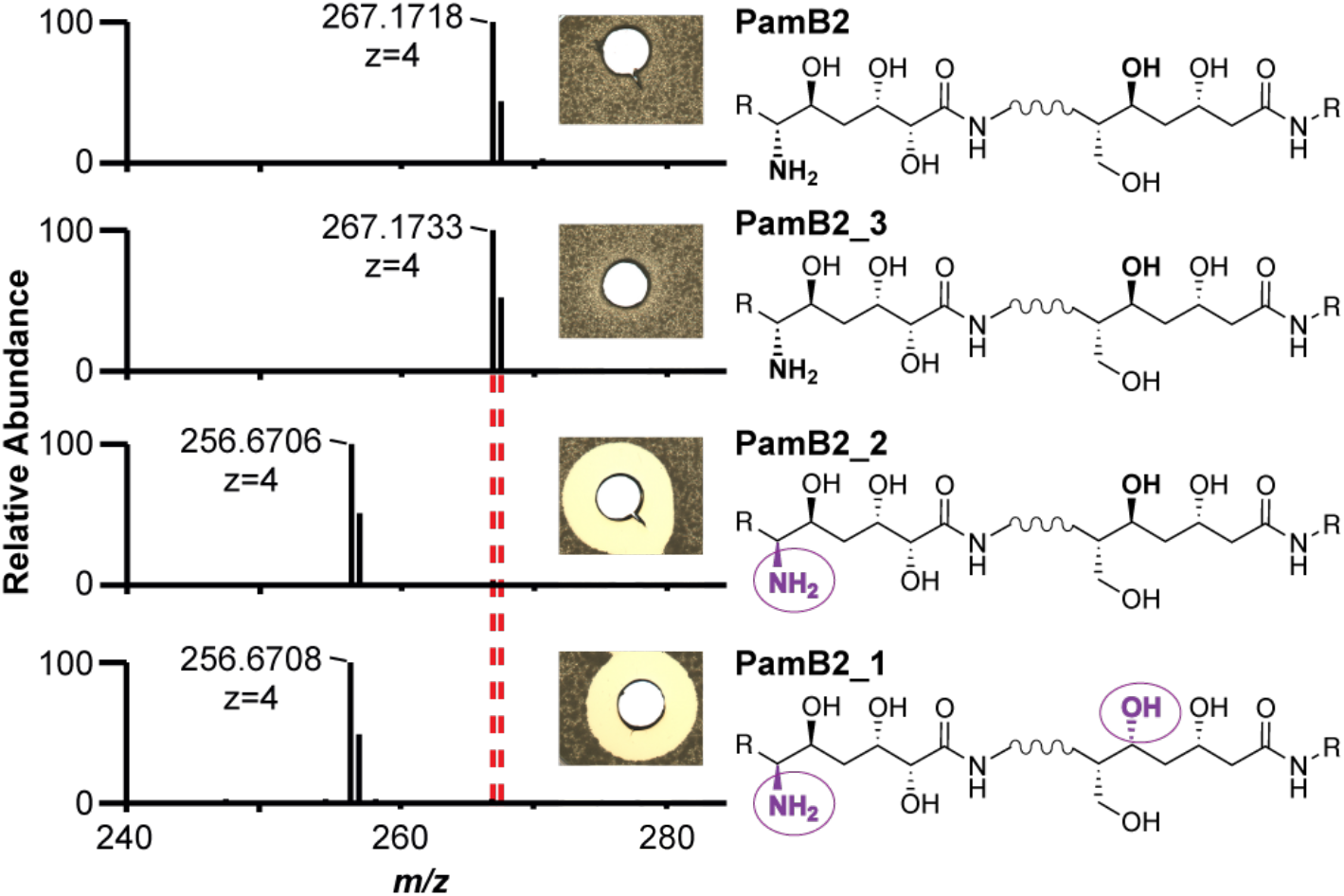
Substrate specificity and stereoselectivity of PamZ. The natural product (PamB2), synthetic paenilamicin B2 (PamB2_3) and synthetic diastereomers of paenilamicin B2 (PamB2_2, PamB2_1) were incubated with PamZ and acetyl-CoA *in vitro* and tested in an agar diffusion assay against *Bacillus megaterium* (insets). Each single reaction was verified by HPLC-ESI MS. Dashed lines indicate the mass shift of 42 Da (4x 10.5 Da) due to *N*-acetylation. Changes in stereoconfiguration are highlighted in purple and circles.

### The structure of PamZ:acetyl-CoA binary complex

A BLAST^29^ search indicated that PamZ belongs to the GNAT superfamily with a sequence identity of 31% to the *N*-acetyltransferase, ZmaR, whose structure has not yet been determined and which confers resistance against the aminopolyol peptide antibiotic, zwittermicin A, in *Bacillus cereus* UW85 (**Figure S24**)^30^. We elucidated the crystal structure of PamZ in complex with acetyl-CoA at a resolution of 1.34 Å by using the uncharacterized *N*-acetyltransferase from *Streptococcus suis* 89/1591 (PDB-ID: 3g3s) for molecular replacement (**Table S3**). The electron density was of excellent quality, allowed the modeling of the entire poly-peptide chain and unambiguously revealed the bound acetyl-CoA (**Figure S25**). PamZ comprises an N-terminal domain (NTD, residues 1-128, secondary structure elements indicated by primes) and a C-terminal domain (CTD, residues 140-275) which both adopt the characteristic GNAT fold (**Figure 5a**)^31^. The two tandem-GNAT domains, that may have originated from a gene duplication event, share low sequence identity (< 20%) and are connected by an α-helical linker (α_bridge_, residues 129-139). The overall fold of each domain is very similar to that of bacterial aminoglycoside *N*-acetyltransferases (AACs), as pairwise structural alignments with several AACs (PDB-IDs: 1bo4, 1m4i, 1s3z) gave root-mean square deviations (RMSDs) of 2.9-4.2 Å for both the NTD and CTD (**Figure S26**)^32^. A structural superimposition between the NTD and CTD of PamZ yielded an RMSD of 4.2 Å for 75 pairs of C_α_ atoms (**Figure S27**)^32^.

**Figure 5.**
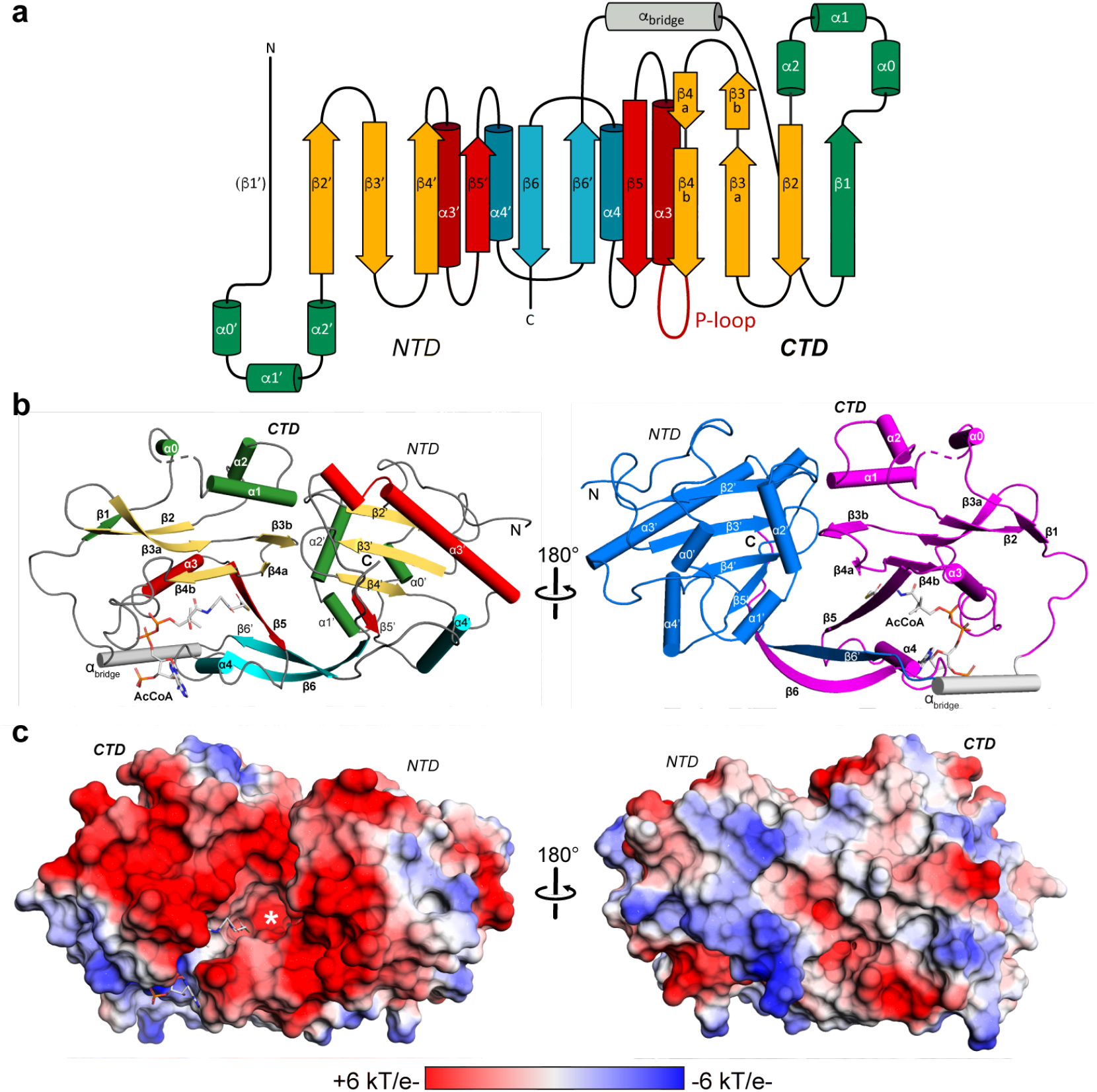
X-ray crystal structure of Gcn5-related *N*-acetyltransferase PamZ. **a** Structural topology of PamZ with its characteristic tandem-GNAT fold. The protein structure is divided into an N-terminal (NTD) and a C-terminal (CTD) domain. Color coding of protein regions follows that of other bacterial GNATs, such as aminoglycoside *N*-acetyltransferases (AACs)^31^. **b** Cartoon representations of PamZ from two perspectives. The first perspective (*left*) follows the color code as in a). Acetyl-CoA (AcCoA) bound to the P-loop and β-bulge of the CTD is depicted as sticks. The β-bulge is formed by strands β4b and β5. The tandem-GNAT domains are highlighted (*right*) in blue (NTD) and purple (CTD). Secondary structure elements are labeled according to the protein topology. **c** Identical view as in panel b) with the electrostatic potential mapped on the surface of PamZ, illustrating positive (blue) and negative (red) charges. The acetyl group attached to CoA (sticks) points into the active site highlighted by an asterisk.

However, a comparison with the typical GNAT fold revealed several unique features in PamZ. Instead of two N-terminal α-helices, α1 and α2, both domains of PamZ contain three short helical segments, α0-α1-α2, which pack onto one face of the central antiparallel β-sheet, β2-β3-β4, whereas helix α3 buries its other side. A kink in the backbone conformation of strand β3, involving residues T199 and C200, causes a strong right twist and thus a distortion of the antiparallel β3-β4 arrangement, which led us to discriminate these strands as β3a/β3b and β4a/β4b (**Figure 5a**). The central β-sheet is extended by strand β5’ in the NTD, whereas the CTD shows the characteristic β-bulge of GNAT enzymes – a V-shaped cavity between strands β4b and β5 accommodates the pantetheine segment of CoA (**Figure 6a**). Furthermore, the well-conserved pyrophosphate-binding loop (P-loop) of the GNAT family (R/Q-X-X-G-X-A/G)^25^ is only present in the CTD of PamZ (Q-N-K-G-L-A) between strand β4b and helix α3 (**Figure 6a**)^33^, whereas the NTD is missing this signature motif. Accordingly, there is only one acetyl-CoA molecule canonically bound in the PamZ structure, namely to the CTD.

**Figure 6.**
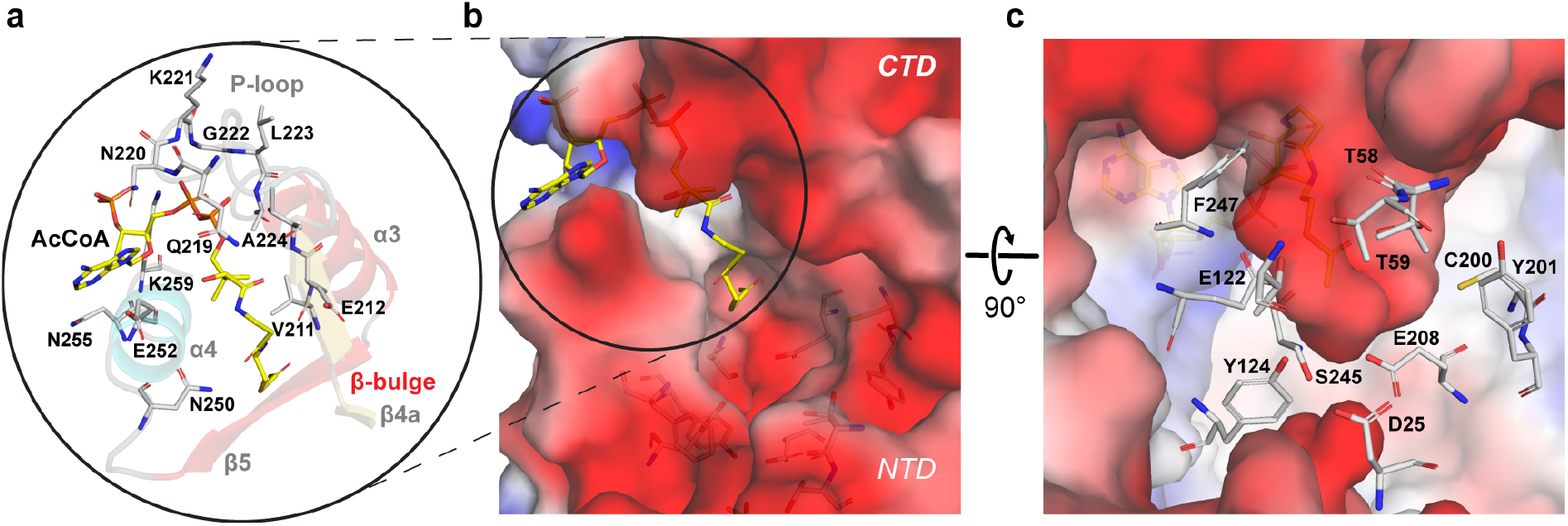
Active site of PamZ. **a** Motifs A (β4-α3) and B (β5-α4) located in the C-terminal domain (CTD) interact with co-substrate acetyl-CoA. **b** Close-up view of the active site displaying the negatively charged groove (color code as in **Figure 5c**). **c** Highlighted amino acid residues with hydrogen-donating and -accepting groups form the groove and interact with the substrate paenilamicin.

Hence, we concluded that the NTD is incompetent of binding acetyl-CoA and rather plays a structural role, in particular for substrate binding (see below). Notably, many GNAT enzymes exist as homodimers in solution with various arrangements of the monomer-monomer interface^31^. Likewise, AACs have often been crystallographically observed in a homodimeric state, although their quaternary structure in solution may vary^34^. PamZ exists as a monomer, both in solution and in the crystal (**Figure S28**). However, the tandem-GNAT domain constellation of PamZ achieves an intramolecular domain-domain interface that resembles that of some GNAT homodimers. There are several GNAT enzymes that utilize domain swapping of strand β6 to stabilize their homodimeric structure^33,35,36^. Interestingly, a major interface in PamZ is achieved by domain-swapping of strand β6 (β6’), which inserts between strands β5’ and β6’ (β5 and β6) of the opposing domain and thus forms an extended, antiparallel and strongly-twisted β-sheet throughout the enzyme (**Figure 5b**). This β-sheet is only interrupted by the β-bulge in the CTD accommodating the cofactor and allowing the amide groups of its pantetheine portion to form pseudo-β-sheet hydrogen-bonds to strand β4b (**Figure 6a**). A very similar tandem arrangement of a pseudo-GNAT NTD and a canonical GNAT CTD can be found in the template protein (PDB-ID: 3g3s). Another example is the structure of mycothiol synthase MshD from *Mycobacterium tuberculosis*, which is also organized as a tandem repeat of two GNAT domains with a catalytically inactive NTD^37^.

PamZ appears to utilize its NTD to form a well-defined substrate pocket with strands β5 and β6’ representing its floor. A second interface between the NTD and CTD is accomplished through tight packing of helix α2’ onto the small β3b-β4a sheet. Further interactions involve helix α2 of the CTD and the loops between α2’ and β2’ as well as β3’ and β4’ of the NTD. These inter-domain contacts fully cover the central groove that is normally found at the interface of homodimeric structures of GNAT enzymes and restrict substrate entry to the opening that is also used by the cofactor. This remaining cleft between the two domains of PamZ is decorated with several acidic residues (*e*.*g*. E89, E116, E118, D120, D162, D170, D215, E216, E217, E218, E272, E274 and the C-terminus) and thus deploys a large negatively-charged surface to attract its polycationic substrate (**Figure 5c**). A corridor that lies aside and beyond the acetyl group of the cofactor is approximately 7-8 Å deep and 8-9 Å wide with respect to the thioester carbonyl atom. Although we did not obtain crystals of a ternary PamZ-acetyl-CoA-paenilamicin complex, the position of acetyl-CoA, the well-defined shape of the neighboring pocket and our knowledge about the substrate’s N-terminal acetylation site allows us to predict that the Glm/Aga side-chain of paenilamicin very likely penetrates into this pocket. Acidic residues D25 (loop between α1’ and α2’), E122 (β6’), and E208 (β4a) are well-positioned within the pocket to accommodate and stabilize the guanidine group of Aga, as well as to tolerate the Nζ amine of Glm. Other residues that shape the substrate pocket include T58/T59 (loop between β3’ and β4’), T98 (β5’) and Y124 (β6’) of the NTD as well as C200/Y201 (β3b) and S245/F247 (β5) of the CTD (**Figure 6c**). This shows that both domains most likely contribute to substrate recognition. Moreover, the structure of PamZ explains its regioselectivity: if PamZ was to modify *e*.*g*. the terminal amino group of spermidine in paenilamicin, the enzyme would not require such a deep substrate-binding pocket. The architecture of the central groove between the NTD and CTD has evolved to optimally accommodate the N-terminal Glm/Aga building block of paenilamicin, whilst terminal amines such as those of spermidine, ornithine and lysine side-chains would not occupy this binding pocket, as they would experience significantly less binding stabilization.

Such accommodation of Glm/Aga in the substrate pocket would position the N-terminal amino group of Aga-6 close to the thioester carbonyl of the cofactor. An active-site aspartate or glutamate residue commonly acts as a general base to trigger the *N*-acetylation reaction by deprotonation of the amine followed by a nucleophilic attack at the carbonyl of the thioester^34^. In PamZ, the side-chains of E122 (β6’) as well as E208 (β4a) exhibit an interatomic distance of approximately 7 Å to the carbonyl atom of acetyl-CoA and thus might be in close proximity to the N-terminal amino group of Aga-6 (**Figure 6c**). Residue S245 (β5) is sandwiched between E122 and E208, and may mediate deprotonation and/or proton shuttling. Furthermore, we cannot exclude the involvement of water molecules during proton transfer. An oxyanion hole as described for myristoyl-CoA transferase^38^ is not present in PamZ, but the amide proton of V211 (β4b) facilitates hydrogen-bonding to the carbonyl oxygen of the thioester, which would increase the electrophilicity of the carbonyl carbon and stabilize the tetrahedral transition state after nucleophilic attack.

### Self-resistance mechanism of *P. larvae*

The deactivation of paenilamicin through formation of *N*-acetylpaenilamicin by the action of PamZ (**Figure S3-S7**) implicates that the enzyme may confer self-resistance to the producer strain *P. larvae*. To test this hypothesis, we exposed the deletion mutant *P. larvae* Δ*pamZ* to a mixture of paenilamicin A1/B1 in an agar diffusion assay. The mixture, which was purified from *P. larvae* ERIC II, inhibited bacterial growth of the deletion mutant Δ*pamZ*, but not that of the WT strain (**Figure 7a**).

**Figure 7.**
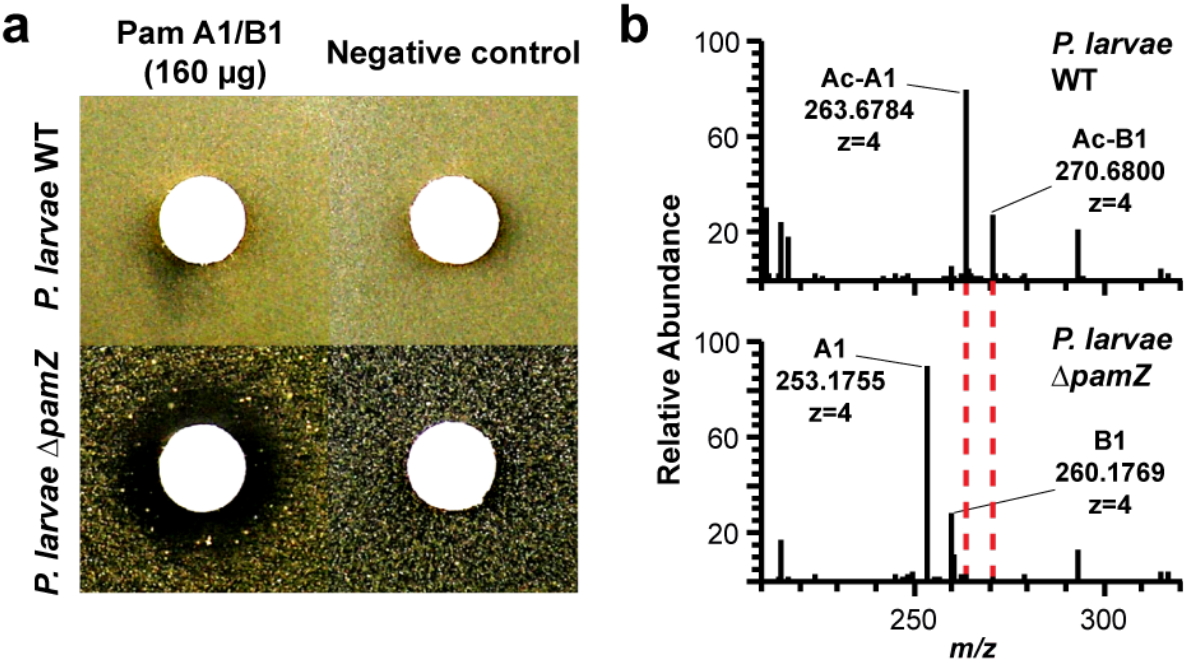
Self-resistance of *P. larvae* against paenilamicin. **a** Deactivation of a paenilamicin mixture A1/B1 (*left*) was tested by an agar diffusion assay against *P. larvae* WT (*top*) and *P. larvae* Δ*pamZ* (*bottom*). The negative control (*right*) contained water only. **b** HPLC-ESI MS spectra of cell lysates of *P. larvae* WT (*top*) and *P. larvae* Δ*pamZ* (*bottom*) are depicted. Relevant peaks for paenilamicin (A1/B1) and *N*-acetylpaenilamicin (Ac-A1/Ac-B1) species are labeled with corresponding *m/z* ratios (*z*=4).

This result demonstrated that *P. larvae* requires the resistance gene, *pamZ*, to protect itself from the deleterious effects of its own antibacterial agent, paenilamicin. For further experimental support, we analyzed supernatants and cell pellets of *P. larvae* WT and Δ*pamZ* for paenilamicins and *N*-acetylpaenilamicins. In cell lysates of *P. larvae* WT, we exclusively found *N*-acetylpaenilamicin, whereas for the deletion mutant Δ*pamZ* only unmodified paenilamicin (**Figure 7b**) was detected. From paenilamicin isolates of the WT strain, primarily paenilamicin and only small amounts of *N*-acetylpaenilamicin were found in the supernatant by HPLC-ESI MS (**Figure S2**). These findings demonstrate that the self-resistance factor PamZ enables *P. larvae* WT to acetylate and thus inactivate intracellular paenilamicin.

*N*-acetylation functions as an efficient self-protection mechanism by scavenging paenilamicin that reenters the cells of *P. larvae*. However, this mechanism may not apply to intracellular paenilamicin after its release from the NRPS-PKS assembly line. Instead, it seems very likely that an inactive precursor, *i*.*e*. a prodrug, of paenilamicin is produced to mask the strong antibacterial activity before cellular export. Along these lines, the biosynthetic gene cluster of paenilamicin^22^ harbors the *pamJ* gene, which shows significant sequence similarity to a cyclic-peptide export ABC transporter with D-asparagine-specific peptidase activity that has been reported to be involved in a prodrug resistance mechanism in nonribosomal peptide synthesis^39–42^. The peptidase recognizes and cleaves an *N*-acyl-D-asparagine unit of the prodrug. Accordingly, *P. larvae* must have developed a dual self-resistance mechanism against paenilamicin both potentially addressing the N-terminal Glm/Aga region, specifically the N-terminal amino group at Aga-6 position, as modification site. Not only *P. larvae*, but also other bacteria belonging to the Firmicutes refer to a dual self-resistance mechanism associated with NRPS-PKS-derived compounds like amicoumacin^41,43^, zwittermicin^30^ and edeine^44^ (**Figure S29**). In a very recent study, paenilamicin B2 showed an inhibitory effect (IC_50_ of approx. 0.3 µM) on the *E. coli* ribosome *in vitro*, whereas the diastereomer PamB2_2 was approx. 10-fold and the *N*-acetylpaenilamicin B2 approx. 100-fold less active^28^. The modifications of the N-terminal amino group at Aga-6 thus point to the importance of the N-terminal Glm/Aga region as major pharmacophore mediating recognition at the molecular target.

The insights into the pharmacophore region of paenilamicins and the structure of PamZ including its substrate-binding pocket may lead to the development of inhibitors against the self-resistance factor to weaken the bee larvae pathogen. In summary, these results expand our knowledge of the molecular strategies exploited by *P. larvae* to survive in its ecological niche – knowledge that is needed to combat this pathogen and secure health of bee colonies worldwide.

## Methods

### Bacterial strains and culture conditions

The field strain *Paenibacillus larvae* (*P. larvae*) 04-309 (DSM 25430) and the deletion mutant 04-309 Δ*pamZ* were cultivated as follows: bacteria were grown on Columbia sheep blood agar (CSA, Thermo Fisher Scientific Oxoid, Schwerte, Germany) medium plates at 37°C for 2-3 days. A preculture of 2 mL Mueller-Hinton-yeast-phosphate-glucose-pyruvate (MYPGP)^45^ medium was inoculated with a single colony and grown overnight. A 50 mL culture of MYPGP broth was inoculated with the preculture to reach an optical density measured at 600 nm (OD_600_) of 0.001. This main culture was incubated at 30°C for 72 h with gentle shaking (80 rpm). Cultures were centrifuged at 3200 g, 4°C for 30 min and supernatants were stored at −20°C until further use. *Escherichia coli* (*E. coli*) BL21-Gold(DE3) cells were cultivated in Luria-Bertani (LB) medium at 37°C and 180 rpm. The medium was supplemented with kanamycin (50 µg mL^-1^) as antibiotic based on the selection marker of the plasmid after transformation. Indicator strains like *B. megaterium* used for the agar diffusion assay were cultivated in LB medium at 37°C and 180 rpm.

### Deletion mutant generation

The generation of the *pamZ* deletion mutant was realized through a well-established protocol for *P. larvae* using the TargeTron Gene Knockout System (Sigma-Aldrich, Germany) based on group II intron insertion as previously described^10,13,14,23,46^. The *pamZ* gene of *P. larvae* DSM 25430 (GenBank: CP003355.1, range from 1729003 to 1729830) was disrupted via site-specific insertion of a 900 bp-sized bacterial mobile group II intron LI.LtrB from *Lactococcus lactis* at position 118 from the start codon. The intron was previously modified to enable specific insertion at this site identified by a computer algorithm provided by the manufacturer (http://www.sigma-genosys.com/targetron) with primers also identified by the computer algorithm (**Table S4**). After successful cloning and transformation into *P. larvae* DSM 25430, screening for *P. larvae* DSM 25430 deletion mutants with the intron integrated in the *pamZ* gene was done via PCR with *pamZ*-specific primers (**Table S4, Figure S30a**). Growth of the *pamZ* deletion mutant in liquid MYPGP medium was not significantly altered in comparison to the wild type strain (**Figure S30b**, two-way-ANOVA, p=0.6486). In brief, growth curves were obtained as follows. *P. larvae* starting cultures had an optical density at 600 nm (OD_600_) of 0.001 and were covered with mineral oil for anaerobic conditions. Cultures were grown in a 96-well-plate (Greiner Bio-One GmbH, Frickenhausen, Germany) and incubated at 37°C while shaking in a Synergy HT plate reader (BioTek, Bad Friedrichshall, Germany). Measurements of the OD_600_ took place hourly for 48 h. The experiment was repeated three times with three biological replicates with three technical replicates each. Representative results are shown.

### Genomic DNA isolation

Cells of *P. larvae* were picked from CSA plates, resuspended in 50 µL water and incubated at 95°C for 10 min. They were centrifuged at 5000 g for 5 min and the supernatant containing the DNA was stored at –20°C until further use. For gene amplification for cloning procedures, pure DNA was isolated by using the MasterPure™ Gram Positive DNA Purification Kit (Epicentre, Illumina, San Diego, CA, USA) following the manufacturer’s instructions.

### Plasmid construction and transformation

Primers were designed for the amplification of the *pamZ* gene from *P. larvae* DSM 25430 and purchased from Thermo Fisher Scientific (**Table S5**). The gene *pamZ* was cloned into vector pET28a(+) introducing an N-terminal histidine-tag and a TEV site. Reactions were performed in the following conditions: initial denaturation at 95°C for 5 min, followed by 30 cycles (105 s per cycle) at 98°C for 30 s, at 61°C for 30 s, and at 72°C for 45 s, followed by a final extension step at 72°C for 10 min. The amplicons were purified and digested with *Nhe*I and *Xho*I, ligated with the digested pET28a(+) vector, and transformed into *E. coli* BL21-Gold(DE3).

### Heterologous expression and protein purification

Terrific broth (TB) medium was inoculated with an overnight culture of pET28a_pamZ transformed in *E. coli* BL21-Gold(DE3) cells to reach an OD_600_ of 0.1 for the purification of PamZ. The culture was incubated at 37°C and 180 rpm until OD_600_ of 0.8-1.0. Expression was induced by addition of 0.2 mM (f.c.) isopropyl β-D-1-thiogalactopyranoside (IPTG). Cells were further incubated at 160 rpm, 18°C for 20 h. Cells were harvested at 5000 g, 4°C for 30 min and the pellet was resuspended in lysis buffer (500 mM sodium chloride, 50 mM TRIS/HCL pH 8.0, 20 mM imidazole). Then, magnesium chloride, DNase, lysozyme and benzamidine were added into the solution. The cell disruption was performed by the cell homogenizer at 15000 psi (Constant Systems Ltd, United Kingdom). Cell lysate was centrifuged at 50000 g, 4°C for 30 min (Beckman Coulter, Avanti J-26 XP). Supernatant was loaded onto a His-Trap column using an ÄKTA protein purification system (GE Healthcare Life Sciences). The chromatography was run with a two-step gradient started with 100% starting buffer (500 mM sodium chloride, 50 mM TRIS/HCL pH 8.0, 20 mM imidazole) and switched to 50% elution buffer (500 mM sodium chloride, 50 mM TRIS/HCL pH 8.0, 250 mM imidazole) within 10 CV to elute the His_6_-tagged PamZ. A His-Trap crude FF column (GE Healthcare Life Sciences) was used for this purification. Fractions of interest were collected and combined to increase protein concentration. Subsequently, TEV protease (1 mg per 10 mg of protein) was added into the concentrated protein solution and incubated at 4°C for 16 h. The N-terminal, TEV-cleavable His_6_-tag was separated from the untagged PamZ by a second nickel affinity chromatography. A size exclusion chromatography was performed with a HiLoad 16/600 Superdex 75 pg column (GE Healthcare Life Sciences) to remove residual imidazole from the protein sample with buffer solution (150 mM sodium chloride, 20 mM TRIS/HCL pH 8.0). The flow rate was set to 1 mL min^-1^. Fractions of interest were collected again, verified by SDS-PAGE and Coomassie staining, and then concentrated. Protein samples were flash-frozen in liquid nitrogen and stored at −80°C for further applications.

### Analytical size exclusion chromatography

Mixture A and B were used as standards. Mixture A contained aprotinin (3 mg mL^-1^), carbonic anhydrase (3 mg mL^-1^), conalbumin (3 mg mL^-1^) and mixture B ribonuclease (3 mg mL^-1^), ovalbumin (4 mg mL^-1^). The chromatograms of mixture A and B were acquired as references to determine the oligomeric state of PamZ. Untagged PamZ (1.25 mg mL^-1^) was prepared to obtain the best fitted chromatogram. The size exclusion chromatography was run with the ÄKTA protein purification system (GE Healthcare Life Sciences), equipped with Superdex 75 10/300 GL and run with buffer solution (150 mM sodium chloride, 20 mM TRIS/HCL pH 8.0). The flow rate was set to 0.5 mL min^-1^.

### Protein crystallization, structure determination and refinement

For crystallization experiments, PamZ was concentrated to 71 mg mL^-1^. Crystallization was performed in a sitting drop vapor diffusion setup at 293 K. The reservoir solution was composed of 40% (w/v) PEG 3350, 50 mM ammonium sulfate and 100 mM sodium acetate at pH 4.6. Prior to flash cooling, the crystals were cryo-protected in a reservoir solution supplemented with 20% (v/v) glycerol. Diffraction data were collected at beamline 14.2 at BESSY. Diffraction data were processed with XDS (**Table S3**)^47^. The structure was solved by molecular replacement with PHASER^48^ using the N-terminal domain of the PDB-ID 3G3S. Since the C-terminal domain could not be readily placed the model was completed by Arp/wArp^49^. The structure was refined by maximum-likelihood restrained refinement in PHENIX^50,51^. Model building and water picking was performed with COOT^52^. Hydrogen atoms for protein residues and ligands were generated with PHENIX.REDUCE^53^. Model quality was evaluated with MolProbity and the JCSG validation server (JCSG Quality Control Check v3.1)^54^. Figures were prepared using PyMOL (Schroedinger Inc.). Electrostatic potentials were calculated with APBS^55^. Structural alignments have been performed using SSM^32^. Structural homologues were identified with the DALI server^56^. Structural interfaces were analyzed with the PISA server^57,58^.

### Compound isolation (supernatant)

1 L of frozen supernatants of *P. larvae* ATCC 9545 or DSM 25430 cultures were thawed and then incubated with Amberlite XAD16 adsorption beads (1 g of beads per 10 mL culture filtrate, Sigma, St. Louis, MO, USA) and stirred for 16 h at room temperature. Then, the flow through was separated from the beads and a three-step gradient elution applied using 1 L of 10% (v/v) methanol followed by 1 L each of 90% (v/v) methanol and 90% (v/v) methanol plus 0.1% formic acid (f.c.) to finally obtain paenilamicin (and also *N*-acetylpaenilamicin). (*N*-acetyl)paenilamicin-containing fractions were concentrated and purified subsequently by using a Grace HPLC column (GROM-Sil 120 ODS-5-ST, 10 µm, 250 × 20 mm) coupled to an Agilent 1100 HPLC system (Agilent Technologies, Waldbronn, Germany) with a MWD UV detector. The separation was accomplished by a linear gradient elution using water plus 0.1% (v/v) formic acid as solvent A and acetonitrile plus 0.1% (v/v) formic acid as solvent B. The gradient started from 3% (v/v) to 15% (v/v) solvent B for 8 min, followed by 100% (v/v) solvent B for 7 min, and finished with an isocratic gradient of 100% (v/v) solvent B for 3 min. The flow rate was set to 20 mL min^-1^. In the next step, paenilamicin-containing fractions were concentrated, adjusted with trifluoroacetic acid to approximately pH 2.0 to increase separation and purified by an Agilent HPLC column (PLRP-S, 100 Å, 10 µm, 150 × 25 mm) coupled to an Agilent 1100 HPLC system with a MWD UV detector for the separation of the native (*N*-acetyl)paenilamicin variants. (*N*-acetyl)paenilamicin was purified by using an isocratic gradient elution using water plus 0.1% (v/v) trifluoroacetic acid as solvent A and acetonitrile plus 0.1% (v/v) trifluoroacetic acid as solvent B. The isocratic gradient was started with 1% (v/v) solvent B for 8 min, followed by a linear gradient from 1% (v/v) to 95% (v/v) solvent B for 7 min, and finished with an isocratic gradient of 95% (v/v) solvent B for 5 min. The flow rate was set to 20 mL min^-1^. (*N*-acetyl)paenilamicin-containing fractions were dried *in vacuo*, lyophilized to obtain pure compound and verified by HPLC-ESI-MS and ^1^H-NMR spectroscopy.

### Compound extraction (cell pellet)

After cultivation of *P. larvae* DSM 25430 and its deletion mutant Δ*pamZ*, the cells were harvested and the cell pellets resuspended in 50% methanol (1 g per 2 mL solvent). The cells were disrupted by sonication (Branson Sonifier 250) for five cycles (15 s each cycle). In between of each cycle, the cell lysate was incubated on ice for 60 s. The lysate was centrifuged at 5000 g, 15°C for 30 min. The supernatant was analyzed for *N*-acetylpaenilamicin and paenilamicin by HPLC-ESI MS.

### *In vitro* activation assay

A reaction mixture consisted of 0.5 mM paenilamicin, 7.5 µM PamZ, 1 mM acetyl-CoA, 1.5 mM sodium phosphate buffer (pH 7.8). Also, samples were prepared each without enzyme and co-substrate as negative controls. The reaction mixture was incubated at 30°C for 8 h. PamZ was removed by Amicon centrifugal filters (Merck KGaA, Germany) using a 10 kDa molecular weight cut-off filter. Deactivation of paenilamicin was tested against *B. megaterium* as indicator strain by agar diffusion assay and analyzed with HPLC-ESI MS and NMR.

For the NMR experiment, excessive acetyl-CoA from the *in vitro* activation assay was removed by using an HPLC column (Phenomenex, Luna C18[2], 100 Å, 5 µm, 100 × 4.6 mm) coupled to an Agilent 1100 HPLC system (Agilent Technologies, Waldbronn, Germany) with a MWD UV detector. The separation was accomplished by a linear gradient elution using water plus 0.1% (v/v) formic acid as solvent A and acetonitrile plus 0.1% (v/v) formic acid as solvent B. The gradient started from 3% (v/v) to 15% (v/v) solvent B for 8 min, followed by 100% (v/v) solvent B for 2 min, and finished with an isocratic gradient of 100% (v/v) solvent B for 2 min. The flow rate was set to 0.6 mL min^-1^.

For the determination of substrate specificity and stereoselectivity of PamZ including synthetic diastereomers, a reaction mixture consisted of 0.5 mM paenilamicin B2 (also for diastereomers), 7.5 µM PamZ, 1 mM acetyl-CoA, 1.5 mM sodium phosphate buffer (pH 7.8). Also, samples were prepared each without enzyme and co-substrate as negative controls. The reaction mixture was incubated at 30°C for 2 h. PamZ was removed by Amicon centrifugal filters (Merck KGaA, Germany) using a 10 kDa molecular weight cut-off filter. After removal of the protein, the reaction mixture was tested against *B. megaterium* as indicator strain by agar diffusion assay and analyzed with HPLC-ESI MS.

### Agar diffusion assay

20 mL of LB medium including 0.75% (w/v) agar was inoculated with bacterial suspension of *B. megaterium* with a final OD_600_ of 0.05. After solidification of the agar plate, holes were punched into the agar for activity testing. 10 µL of each sample was pipetted into the holes after removal of PamZ and the plate incubated at 37°C overnight.

### *In vivo* activation assay

The growth of wild type *P. larvae* DSM 25430 was compared to the growth of *P. larvae* DSM 25430 Δ*pamZ* in the presence of purified paenilamicin A1/B1 from bacteria supernatants in an agar diffusion assay. In brief, pre-cultures with 5 mL volume were grown in MYPGP broth at 37°C while gently shaking overnight. Liquid lukewarm MYPGP agar was inoculated with *P. larvae* pre-cultures to result in a final optical density OD_600_ of 0.05. Agar plates were poured and let harden. Meanwhile, 20 µL of paenilamicin A1/B1 dissolved in MilliQ (in total 160 µg per disk) were dispensed on filter disks and dried at room temperature. The dry filter disks were placed on the agar. The agar plates were incubated at 37°C overnight. Clear zones of inhibition around the filter disks were indicative of a loss of paenilamicin resistance.

### Mass spectrometry analysis

A 6530 Accurate-Mass Quadrupole Time-of-Flight (Q-TOF) LC/MS (Agilent Technologies, Waldbronn, Germany) was used to verify (*N*-acetyl)paenilamicin-containg fractions during the isolation and purification of paenilamicin, The Q-TOF was attached to an Agilent 1260 Infinity HPLC system and equipped with a HPLC column (Poroshell 120, EC-C8, 2.7 µm, 2.1 × 50 mm, Agilent Technologies, Waldbronn, Germany). The HPLC was started with a linear gradient from 5% (v/v) to 100% solvent B for 10 min using water plus 0.1% (v/v) formic acid as solvent A and acetonitrile plus 0.1% (v/v) formic acid as solvent B, followed by an isocratic gradient of 100% (v/v) for 1 min. The column was equilibrated with 5% (v/v) solvent B for 3 min. The flow rate was set to 0.5 mL min^-1^. Other parameters were set as follows: positive mode, gas temperature to 300°C, drying gas to 8 L min^-1^, nebulizer to 35 psi, sheath gas temperature to 350°C, sheath gas flow to 11 L min^-1^, capillary voltage to 3500 V, fragmentor to 330 V, skimmer to 65 V, acquired rate to 1 spec s^-1^.

A LTQ-Orbitrap XL hybrid ion trap-orbitrap (Thermo Fisher Scientific GmbH, Bremen, Germany) was used to verify the *in vitro* activation assays and to generate tandem mass spectra of paenilamicin and *N*-acetylpaenilamicin in data-dependent acquisition (DDA) mode. The LTQ-Orbitrap XL was attached to an analytical HPLC 1200 Infinity system (Agilent Technologies, Waldbronn, Germany) and equipped with a HPLC column (Poroshell 120, EC-C18, 2.7 µm, 2.1 × 50 mm, Agilent Technologies, Waldbronn, Germany). HPLC was run with a linear gradient using water plus 0.1% (v/v) formic acid as solvent A and acetonitrile plus 0.1% (v/v) formic acid as solvent B from 5% (v/v) to 100% (v/v) solvent B for 6 min, followed by an isocratic gradient of 100% (v/v) solvent B for 2 min. The column was equilibrated with 5% (v/v) solvent B for 2 min. The flow rate was set to 0.5 mL min^-1^. The ESI source parameters were set as follows: product ion spectra were recorded in data-dependent acquisition (DDA) mode with a mass range from *m/z* 180 to *m/z* 2000 (MS1: FTMS, normal, 60000, full, positive. MS2: FTMS, normal, 30000). The parameter for the DDA mode was set as follows: dynamic exclusion (repeat count: 3, repeat duration: 30 s, exclusion size list: 50, exclusion duration: 180 s), current scan event (minimum signal threshold: 10000), activation (type: CID, default charge state: 2, isolation width: *m/z* 2.0, normalized collision energy: 35, activation Q: 0.25, activation time: 30 ms).

### Nuclear magnetic resonance spectroscopy

NMR experiments were performed on a Bruker Avance III 700 MHz spectrometer equipped with a room-temperature TXI probe (Bruker, Karlsruhe, Germany). TopSpin 3.5 (Bruker, Karlsruhe, Germany) was used for data acquisition and processing. Spectra analysis was performed using NMRFAM-SPARKY^59–61^. ^1^H and ^1^H-^13^C HSQC spectra of paenilamicin and *N*-acetylpaenilamicin were recorded using samples in D_2_O with 0.1% acetic acid-d4 at 298 K. ^1^H-^13^C HSQC spectra were recorded with acquisition times of 120 ms and 9 ms in the direct ^1^H and indirect ^13^C dimension, respectively. A delay Δ/2 of 1.72 ms was used for INEPT transfers corresponding to ^1^*J*_HC_ of 145 Hz. Apodization of time domain data was performed using a squared sine-bell function shifted by 90°. The 2D data was processed by applying linear forward prediction in the indirect ^13^C dimension and zero filling prior to Fourier transformation. ^1^H chemical shifts were referenced externally using a sample of trimethylsilylpropanoic acid (TMSP-*d*_4_, Deutero GmbH, Kastellaun, Germany) in D_2_O with 0.1% acetic acid-d4 measured at 298 K. ^13^C chemical shifts were referenced indirectly using a correction factor of *f*_13C/1H_ = 0.251449530^62,63^. Chemical shift perturbations (CSPs) were calculated using the following equation^64^:

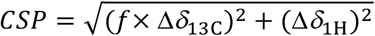

where Δ*δ*_13C_ and Δ*δ*_1H_ correspond to the ^13^C and ^1^H chemical shift differences between paenilamicin B2 and *N*-acetylpaenilamicin B2 for each carbon-proton pair. We used a weighting factor *f* of 0.06 to account for the much larger chemical shift dispersion in the ^13^C dimension (ca. 60 ppm) compared to that in the ^1^H dimension (ca. 3.5 ppm).

## Data Availability

The coordinates and structure factors have been deposited in the Protein Data Bank under accession code 7B3A. Diffraction images have been deposited at www.proteindiffraction.org.

## Acknowledgements

We thank the Deutsche Forschungsgemeinschaft (DFG, German Research Foundation) with SU 239/21-1 (project no. 279410221) and with RTG 2473 “Bioactive Peptides” (project no. 392923329) for funding the project. This research was also funded by the Ministries responsible for Agriculture of the German Federal States of Brandenburg, Sachsen-Anhalt, Thüringen, Sachsen and the Senate of Berlin, Germany, as well as by the DFG, grant numbers GE1365/1-1, GE1365/1-2, and GE1365/2-1 to E.G. We are grateful to Claudia Alings, Freie Universität Berlin, for help with crystallization. We acknowledge access to beamlines of the BESSY II storage ring (Berlin, Germany) via the Joint Berlin MX-Laboratory sponsored by Helmholtz-Zentrum Berlin für Materialien und Energie, Freie Universität Berlin, Humboldt-Universität zu Berlin, Max-Delbrück-Centrum, Leibniz-Institut für Molekulare Pharmakologie and Charité-Universitätsmedizin Berlin.

## Author information

### Contributions

T.D., S.M., R.S., A.M. and R.D.S. designed the experiments. T.D., S.M. and R.S. purified paenilamicins and PamZ and also conducted the *in vitro* activation assays. T.D. set up the tandem-MS experiments and analyzed related data. B.L. performed the crystallization and elucidated the protein structure of PamZ. J.E. generated the deletion mutant *P. larvae* Δ*pamZ* and performed the *in vivo* activation assay of paenilamicin against wild type and deletion mutant. T.B. and S.G. synthesized paenilamicin B2 and the two diastereomers. J.G. cultivated *P. larvae* wild type and deletion mutant Δ*pamZ* and prepared the corresponding supernatants. A.M. performed the NMR experiments, acquired and analyzed the related data. T.D., M.C.W., E.G., A.M. and R.D.S. wrote the manuscript.

## Supplementary Information

Supplementary Figures and Tables.

