## Supplemental File for "Molecular Basis of Antibiotic Self-Resistance in a Bee Larvae Pathogen"

### Table of Contents

**SUPPLEMENTARY FIGURES** **2**
**SUPPLEMENTARY TABLES** **22**
**REFERENCES** **26**

### Supplementary Figures

**Figure S1. Elution profile and corresponding SDS-PAGE PamZ after protein purification including TEV digestion.**

**Figure S2.** Deconvoluted MS$^{1}$ spectra of native paenilamicin variants observed in the supernatant of *P. larvae* a) ATCC 9545 (ERIC I) and b) DSM 25430 (ERIC II) after purification with Amberlite XAD16.

**Figure S3.** MS$^{1}$ spectra of *in vitro* activation assay including paenilamicin A1, acetyl-CoA and PamZ.

**Figure S4.** MS$^{1}$ spectra of *in vitro* activation assay including paenilamicin A2, acetyl-CoA and PamZ.

**Figure S5.** MS$^{1}$ spectra of *in vitro* activation assay including paenilamicin B1, acetyl-CoA and PamZ.

**Figure S6.** MS$^{1}$ spectra of *in vitro* activation assay including paenilamicin B2, acetyl-CoA and PamZ.

**Figure S7.** MS$^{1}$ spectra of *in vitro* time-dependent activation assay including paenilamicin B2 (synthetic), acetyl-CoA and PamZ.

**Figure S8.** MS$^{2}$ spectra of paenilamicin A1 isolated from *P. larvae* DSM 25430 (ERIC II).

**Figure S9.** MS$^{2}$ spectra of paenilamicin A2 isolated from *P. larvae* ATCC 9545 (ERIC I).

**Figure S10.** MS$^{2}$ spectra of paenilamicin B1 isolated from *P. larvae* DSM 25430 (ERIC II).

**Figure S11.** MS$^{2}$ spectra of paenilamicin B2 isolated from *P. larvae* ATCC 9545 (ERIC I).

**Figure S12.** MS$^{2}$ spectra of paenilamicin B2 (synthetic).

**Figure S13.** MS$^{2}$ spectra of *N*-acetylpaenilamicin A1 converted *in vitro* by PamZ.

**Figure S14.** MS$^{2}$ spectra of *N*-acetylpaenilamicin A2 converted *in vitro* by PamZ.

**Figure S15.** MS$^{2}$ spectra of *N*-acetylpaenilamicin B1 converted *in vitro* by PamZ.

**Figure S16.** MS$^{2}$ spectra of *N*-acetylpaenilamicin B2 converted *in vitro* by PamZ.

**Figure S17.** MS$^{2}$ spectra of *N*-acetylpaenilamicin B2 (synthetic) converted *in vitro* by PamZ.

**Figure S18. MS$^{2}$ fragmentation of different paenilamicin and *N*-acetylpaenilamicin variants to determine regioselective acetylation.**

**Figure S19.** MS$^{1}$ spectra of *N*-acetylpaenilamicin A1, B1 and B2 isolated from *P. larvae*.

**Figure S20.** MS$^{2}$ spectra of *N*-acetylpaenilamicin B2 isolated from *P. larvae* DSM 25430 (ERIC II).

**Figure S21.** MS$^{2}$ spectra of *N*-acetylpaenilamicin B1 isolated from *P. larvae* DSM 25430 (ERIC II).

**Figure S22.** MS$^{2}$ spectra of *N*-acetylpaenilamicin A1 isolated from *P. larvae* ATCC 9545 (ERIC I).

**Figure S23. Stereoselective *N*-acetylation of PamZ.**

**Figure S24. Multiple sequence alignment of Gcn5-related *N*-acetyltransferases of PamZ (paenilamicin), ZmaR (zwittermicin) and EdeQ (edeine).**

**Figure S25.** Polder electron density map of acetyl-CoA.

**Figure S26. Structural comparison of PamZ and AACs.**

**Figure S27. Pairwise structural alignment between NTD and CTD of PamZ.**

**Figure S28. Size exclusion chromatogram and calibration curve of PamZ including mixture A and B as calibration standards detected at 280 nm.**

**Figure S29. Biosynthetic gene cluster and *N*-acetylation of paenilamicin, zwittermicin A, edeine and amicoumacin.**

**Figure S30. Determination of successful intron insertion into the *pamZ* gene of *P. larvae* DSM 25430.**

**Figure S31.** $^{1}$H-NMR spectrum of paenilamicin A2 isolated from *P. larvae* ATCC 9545 (ERIC I).

**Figure S32.** $^{1}$H-NMR spectrum of paenilamicin B2 isolated from *P. larvae* ATCC 9545 (ERIC I).

**Figure S33.** $^{1}$H-NMR spectrum of paenilamicin A1 isolated from *P. larvae* DSM 25430 (ERIC II).
**Figure S34.** $^{1}$H-NMR spectrum of paenilamicin B1 isolated from *P. larvae* DSM 25430 (ERIC II).

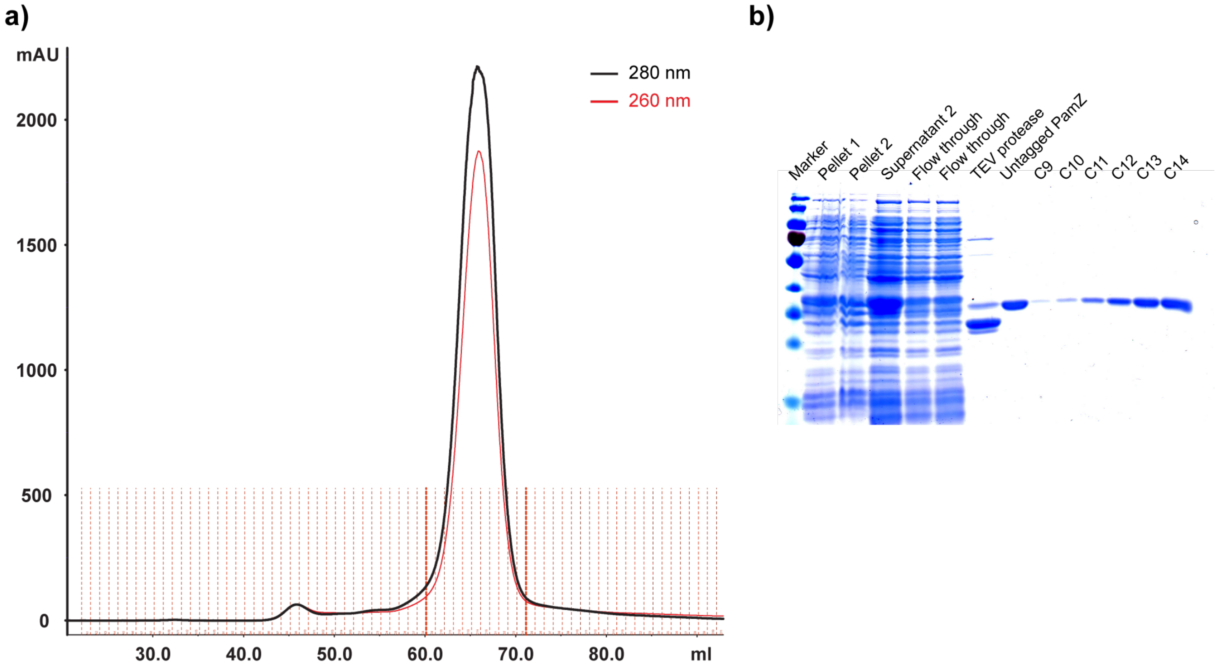


**Figure S1. Protein purification of PamZ including TEV cleavage.** a) Elution profile of size exclusion chromatography after TEV cleavage observed at 280 nm. b) SDS-PAGE of PamZ after protein purification. Fractions C9-C14 were obtained from size exclusion chromatography and concentrated for further applications.

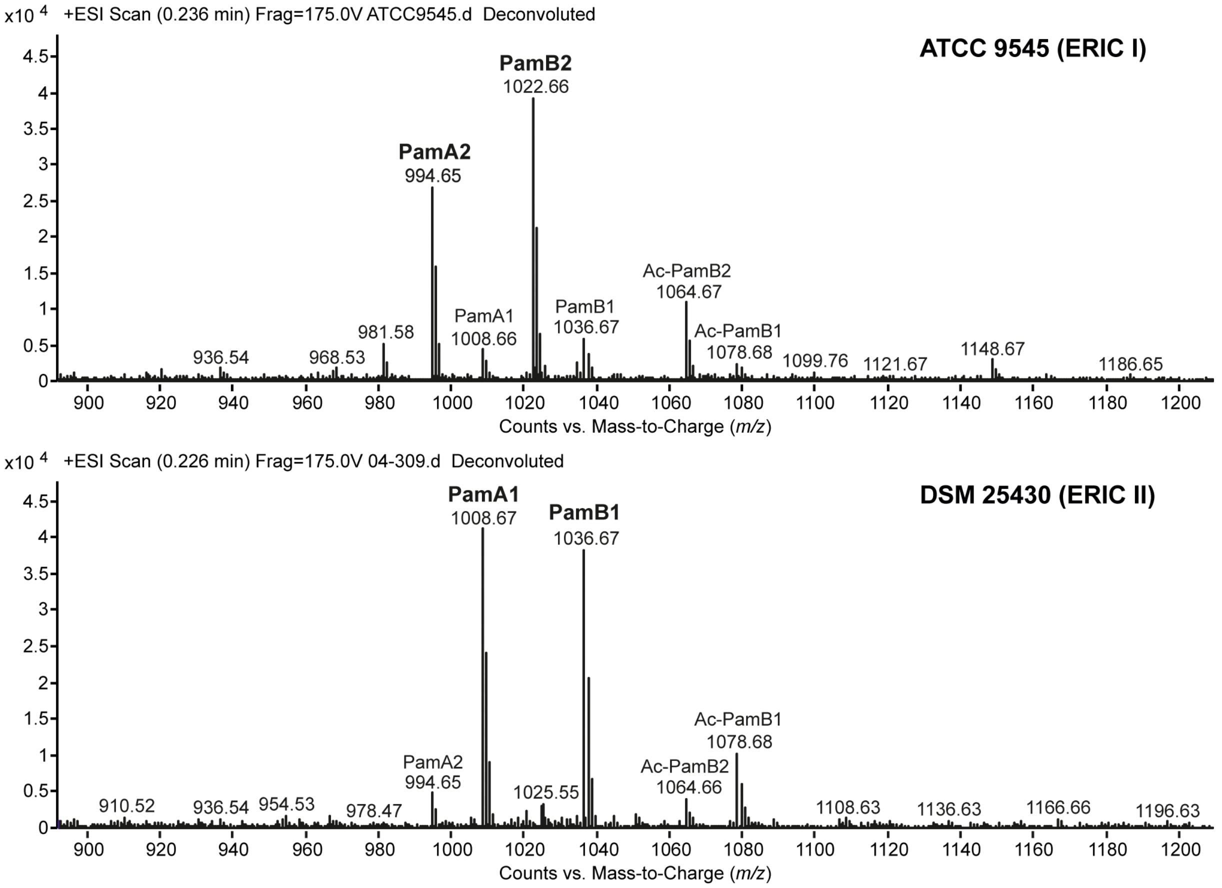


**Figure S2.** Deconvoluted $MS^1$ spectra of native paenilamicin variants observed in the supernatant of *P. larvae* ATCC 9545 (ERIC I) and DSM 25430 (ERIC II) after purification with Amberlite XAD16.

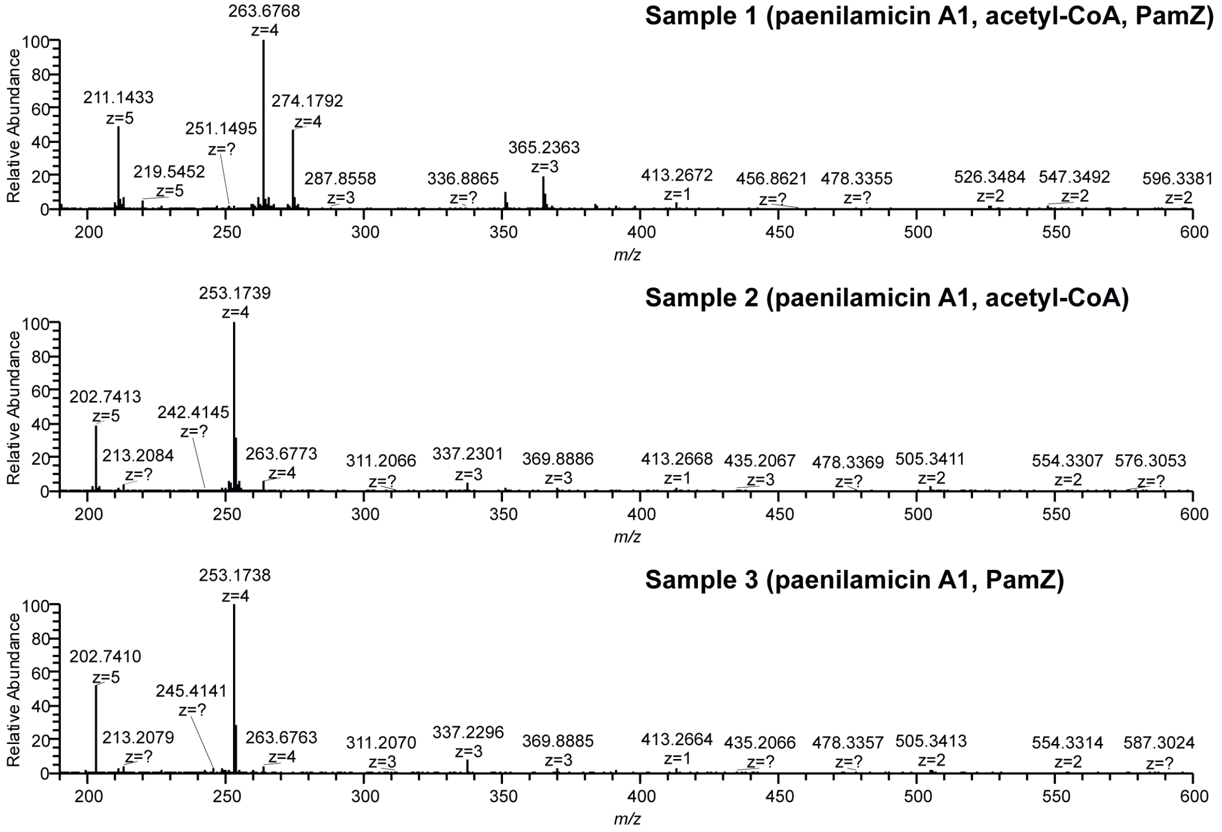


**Figure S3.** MS[1] spectra of *in vitro* activation assay including paenilamicin A1, acetyl-CoA and PamZ.

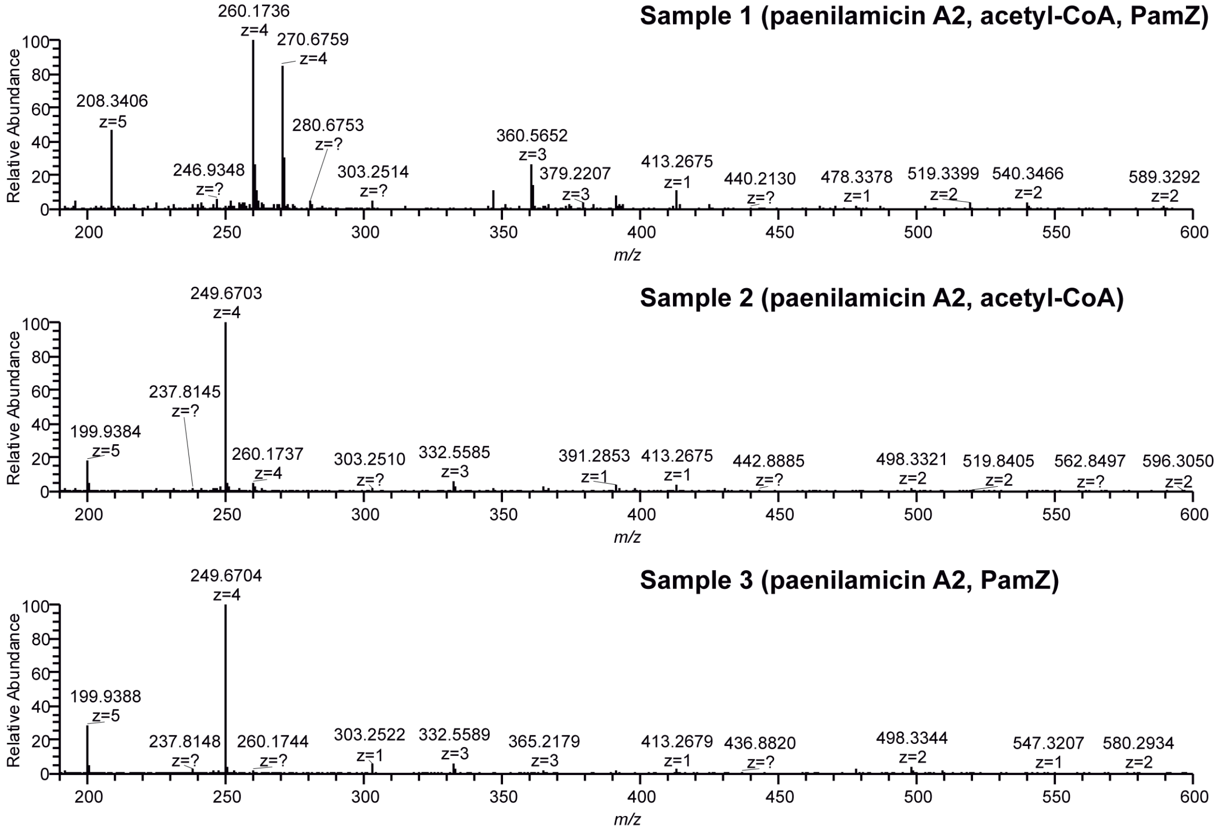


**Figure S4.** MS[1] spectra of *in vitro* activation assay including paenilamicin A2, acetyl-CoA and PamZ.

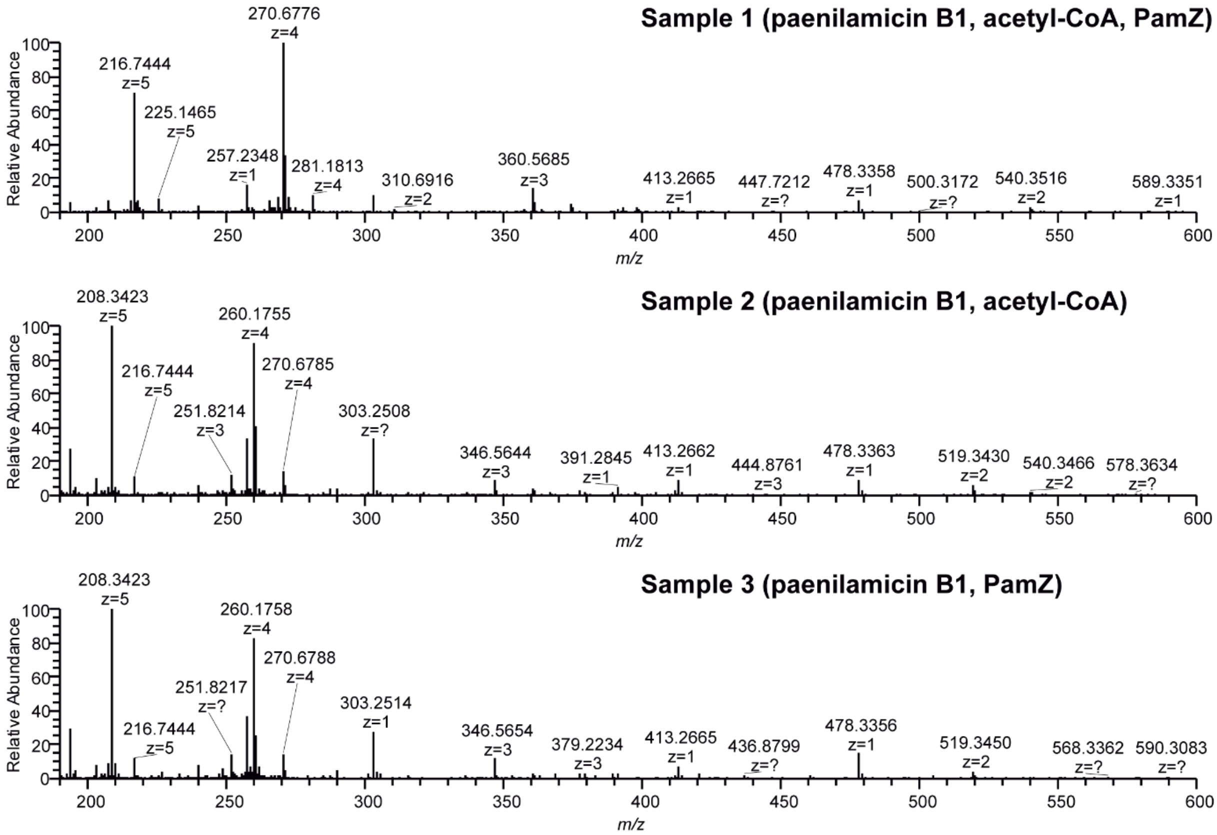


**Figure S5.** $MS^1$ spectra of *in vitro* activation assay including paenilamicin B1, acetyl-CoA and PamZ.

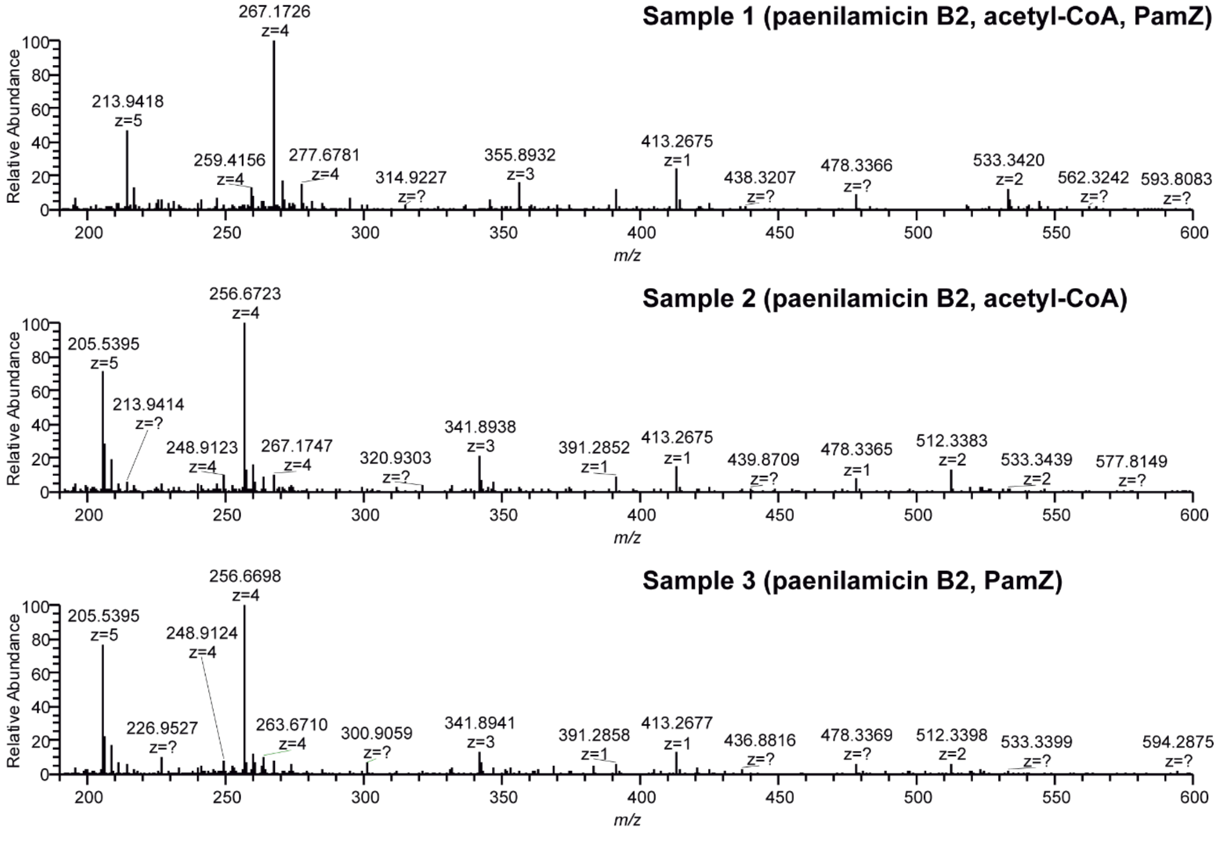


**Figure S6.** $MS^1$ spectra of *in vitro* activation assay including paenilamicin B2, acetyl-CoA and PamZ.

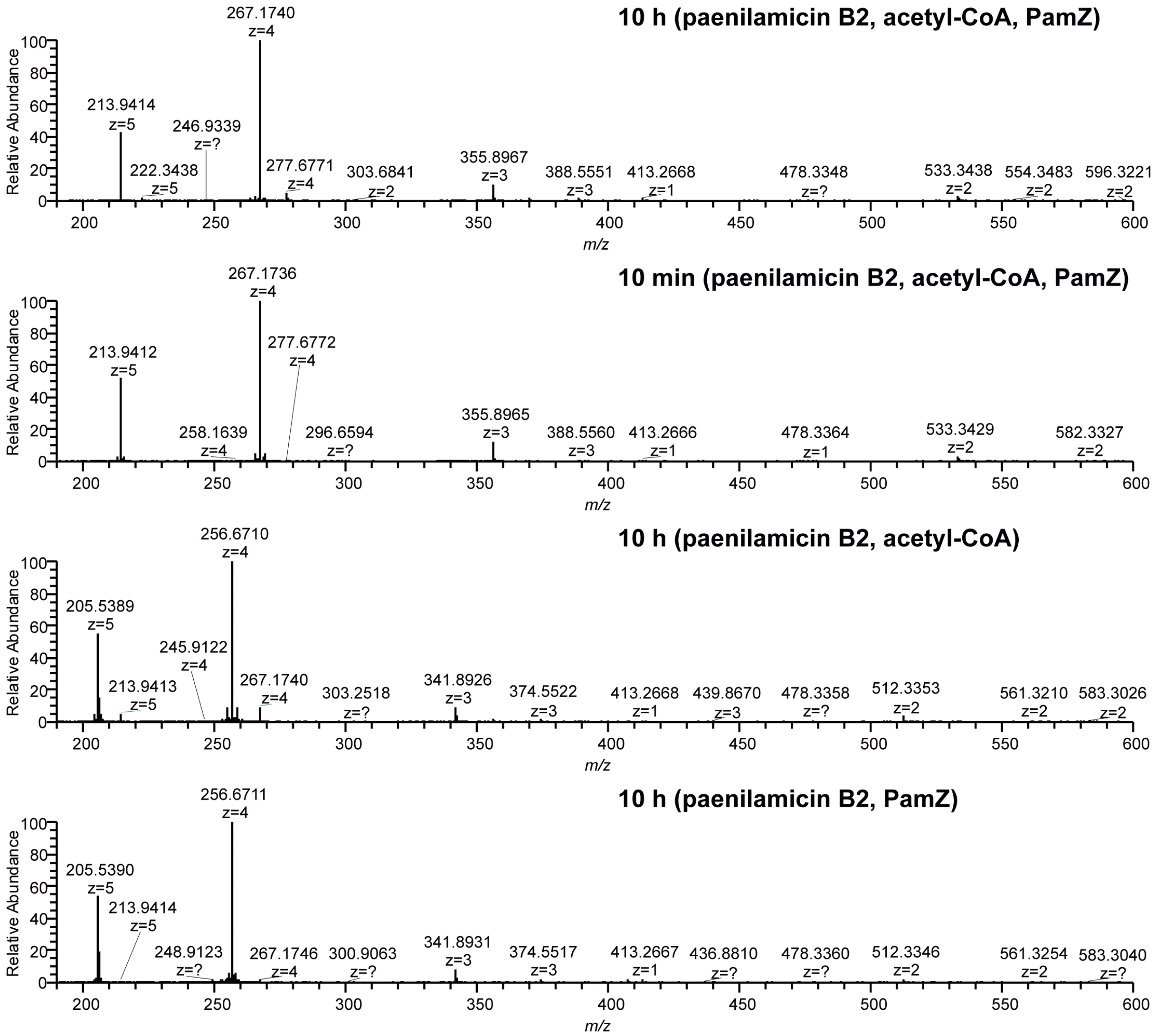


**Figure S7.** $MS^1$ spectra of *in vitro* time-dependent activation assay including paenilamicin B2 (synthetic), acetyl-CoA and PamZ.

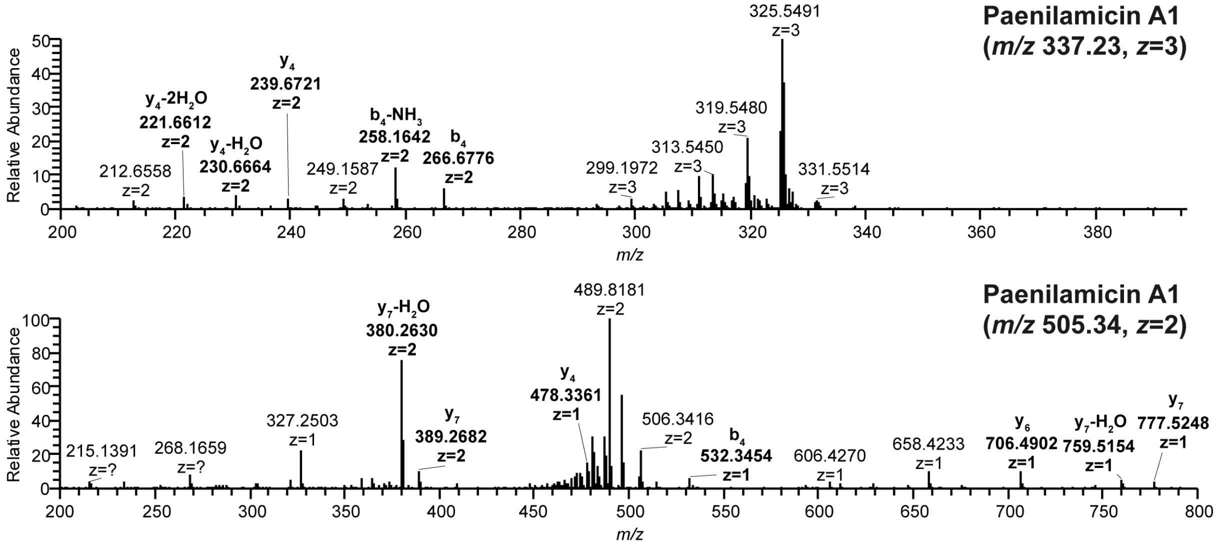


**Figure S8.** $MS^2$ spectra of paenilamicin A1 isolated from *P. larvae* DSM 25430 (ERIC II).

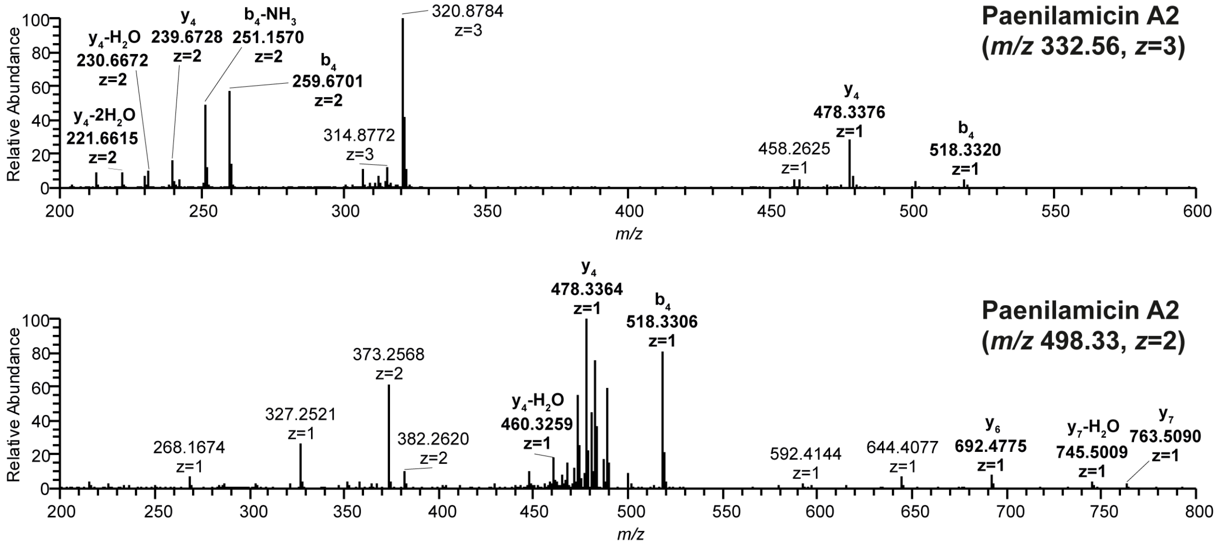


**Figure S9.** $MS^2$ spectra of paenilamicin A2 isolated from *P. larvae* ATCC 9545 (ERIC I).

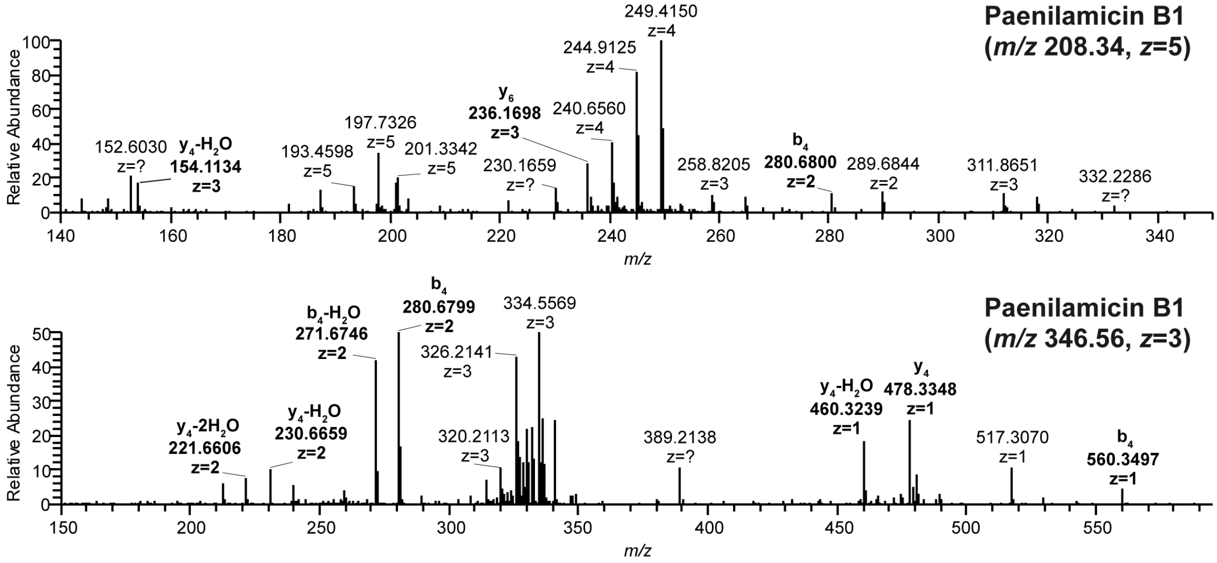


**Figure S10.** $MS^2$ spectra of paenilamicin B1 isolated from *P. larvae* DSM 25430 (ERIC II).

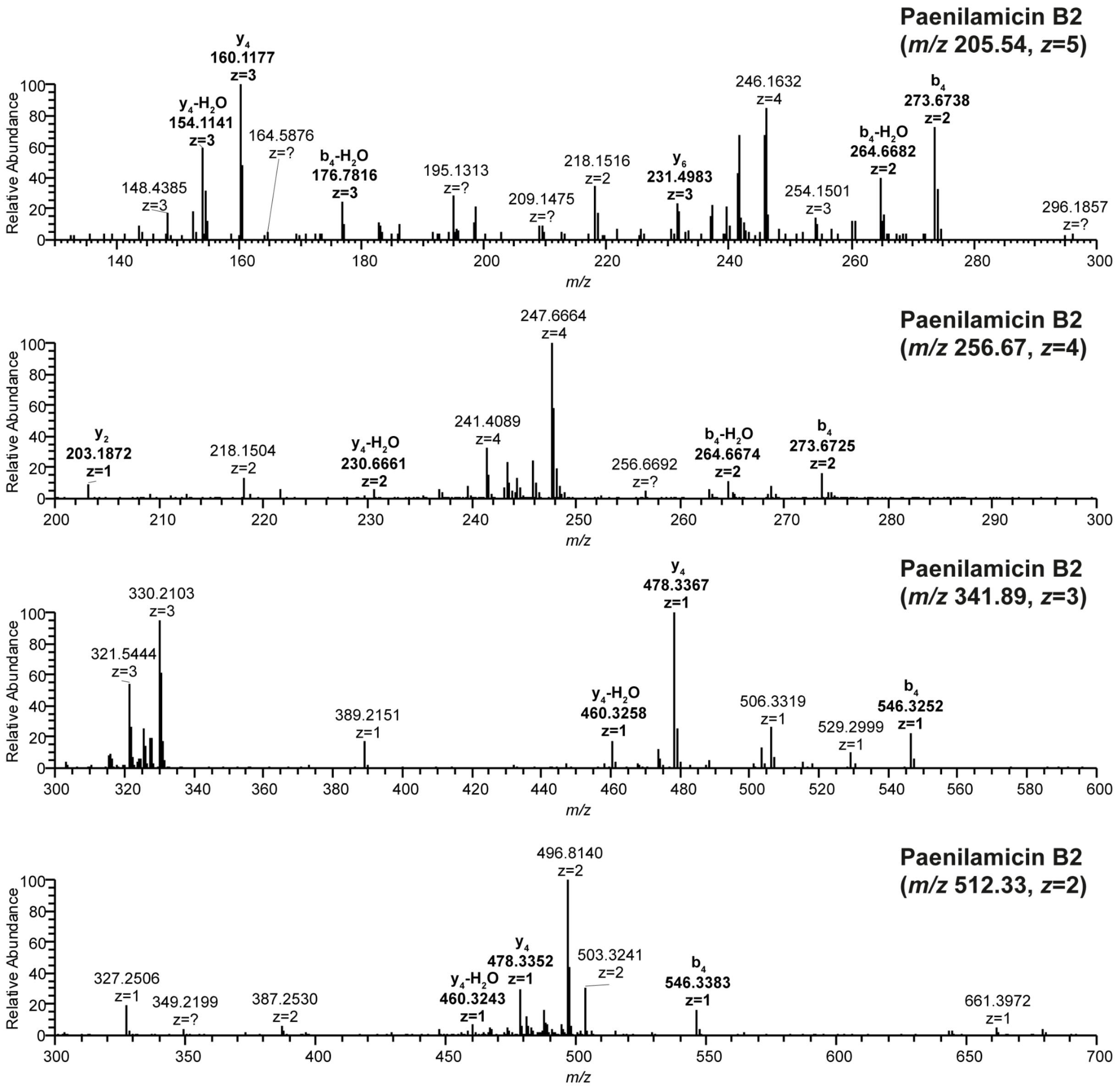


**Figure S11.** $MS^2$ spectra of paenilamicin B2 isolated from *P. larvae* ATCC 9545 (ERIC I).

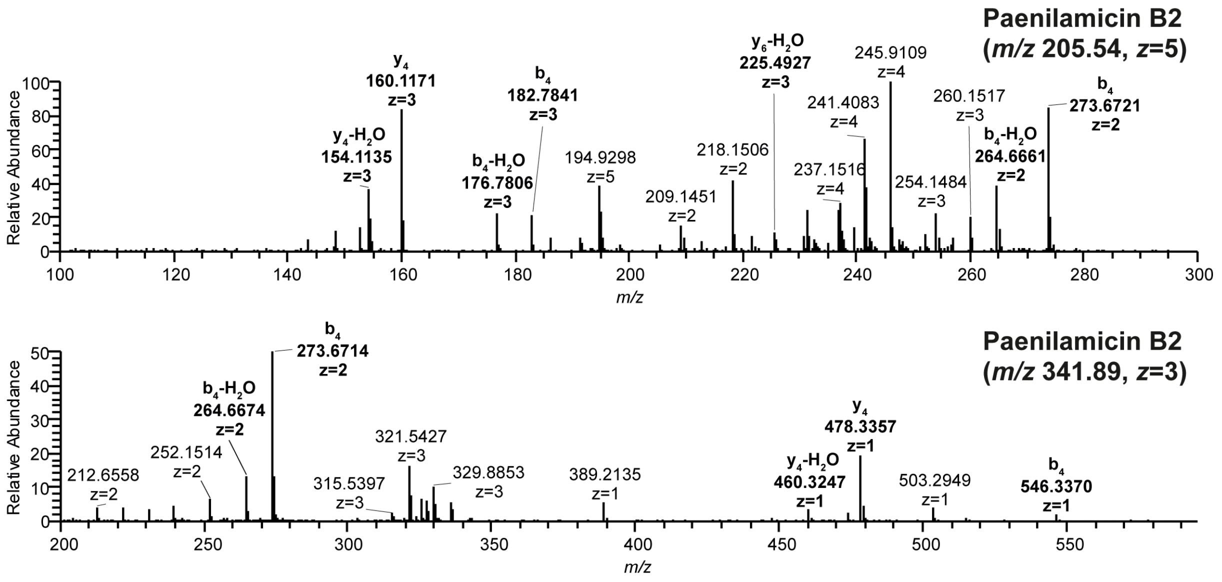


**Figure S12.** $MS^2$ spectra of paenilamicin B2 (synthetic).

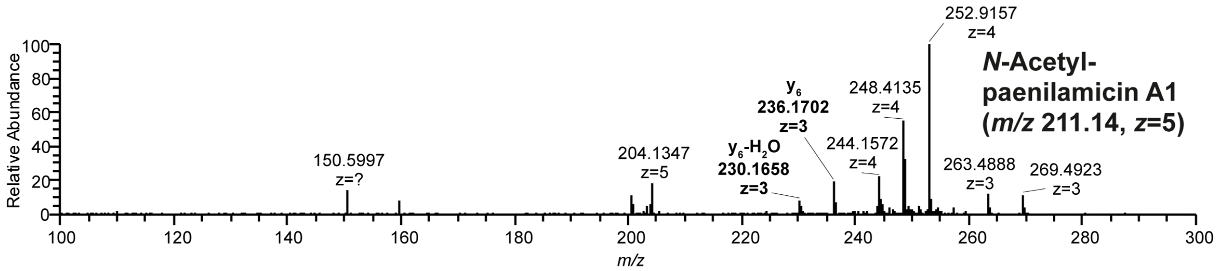


**Figure S13.** $MS^2$ spectra of *N*-acetylpaenilamicin A1 converted *in vitro* by PamZ.

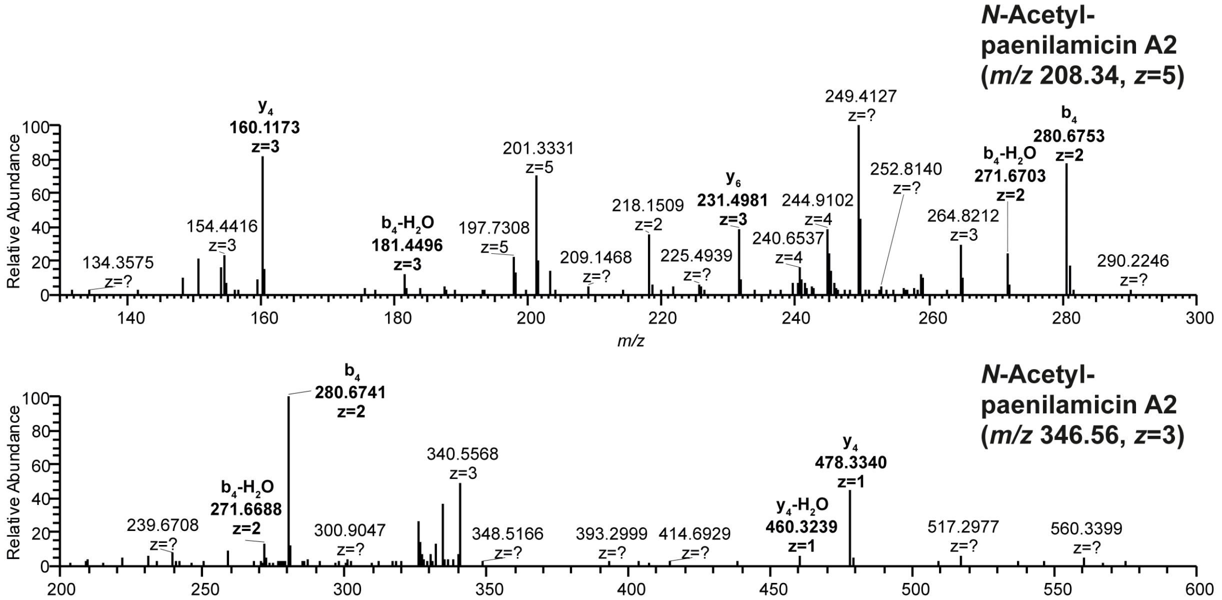


**Figure S14.** $MS^2$ spectra of *N*-acetylpaenilamicin A2 converted *in vitro* by PamZ.

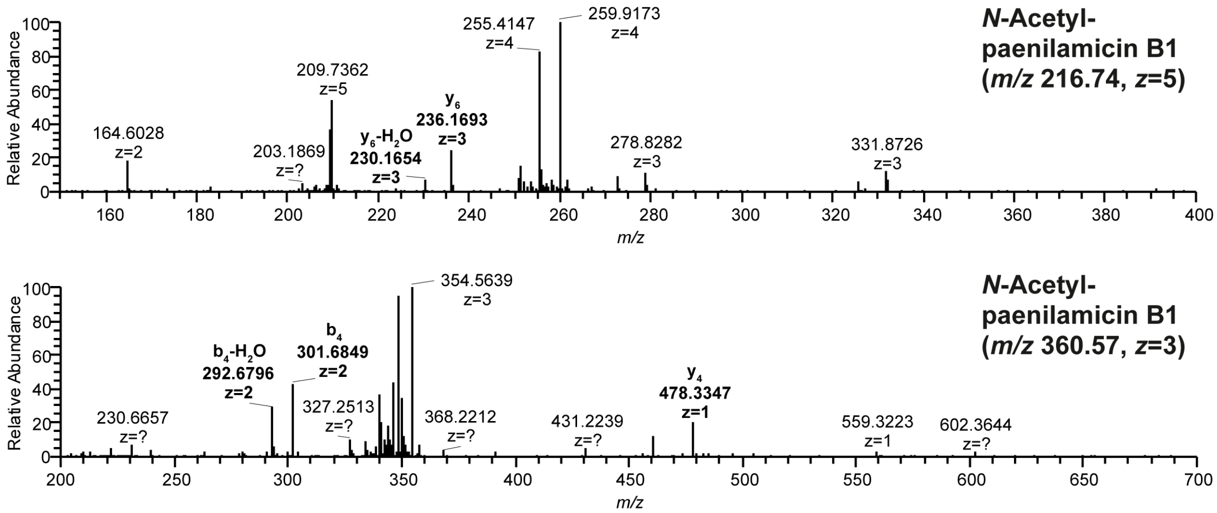


**Figure S15.** $MS^2$ spectra of *N*-acetylpaenilamicin B1 converted *in vitro* by PamZ.

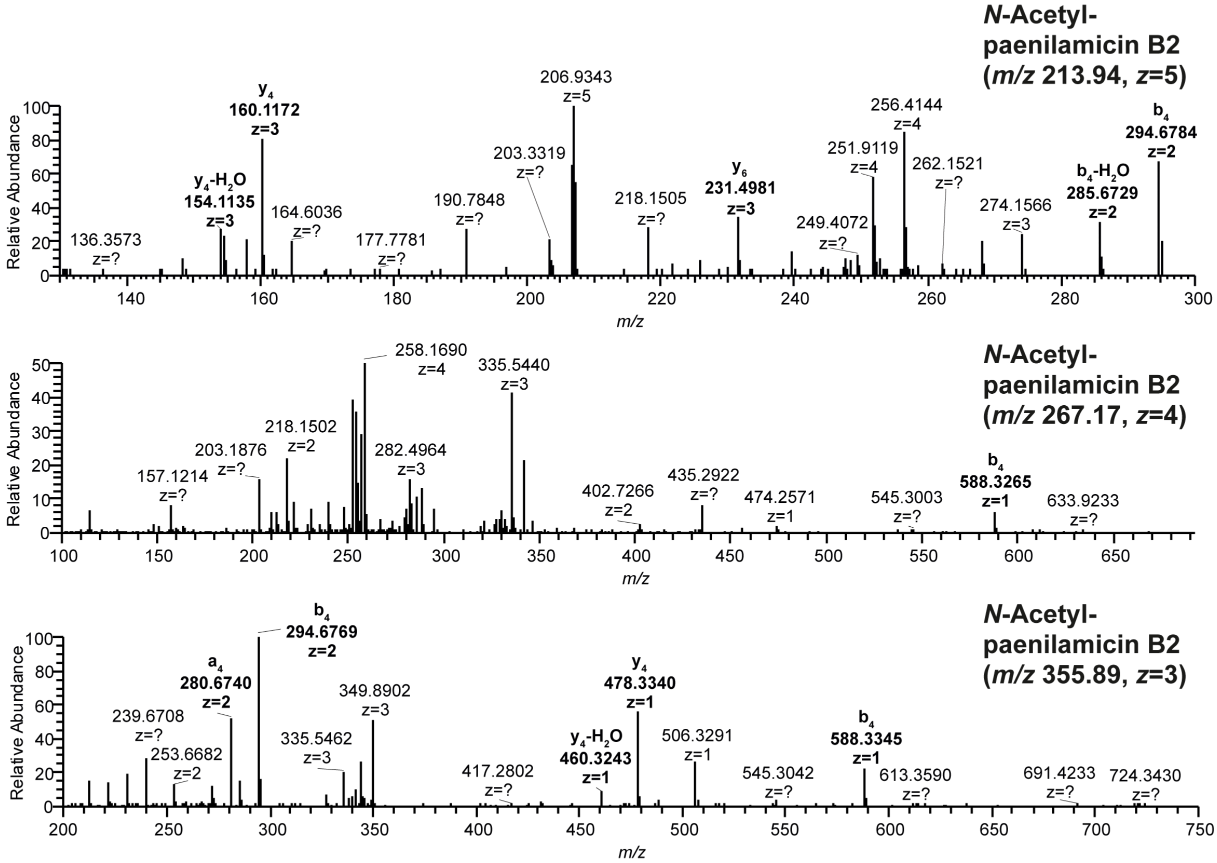


**Figure S16.** $MS^2$ spectra of *N*-acetylpaenilamicin B2 converted *in vitro* by PamZ.

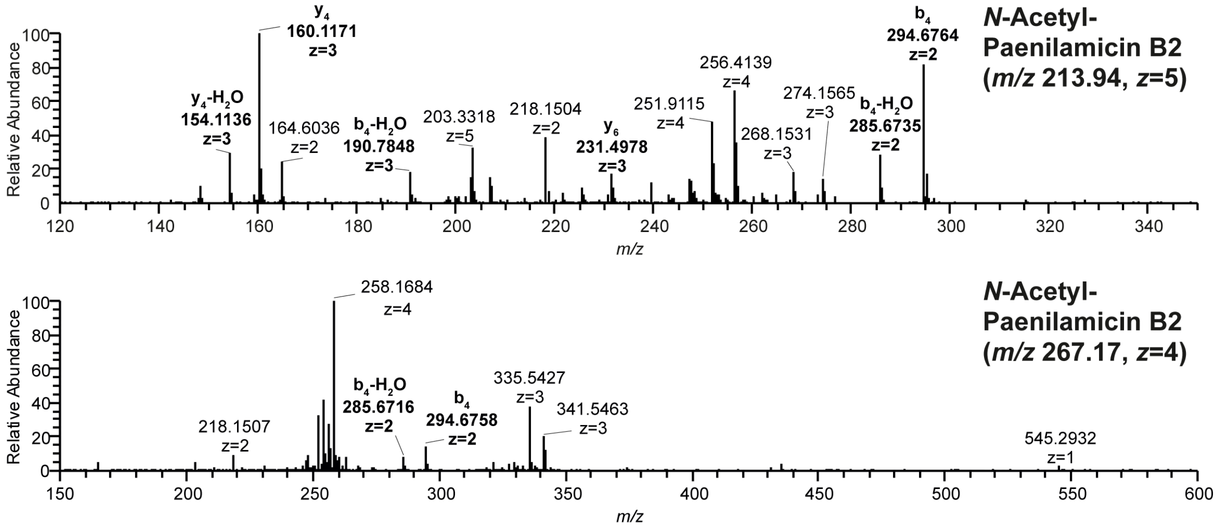


**Figure S17.** $MS^2$ spectra of *N*-acetylpaenilamicin B2 (synthetic) converted *in vitro* by PamZ.

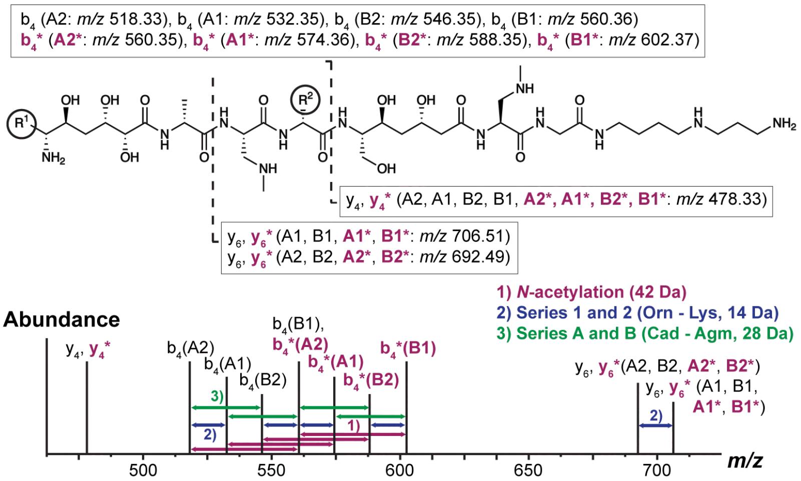


| Derivative | Exact mass | $b_4$ | $y_4$ | $y_6$ |
|---|---|---|---|---|
| A2 | 994.6499 | 518.33 | 478.33 | 692.48 |
| A1 | 1008.6655 | 532.35 | 478.33 | 706.49 |
| B2 | 1022.6560 | 546.35 | 478.33 | 231.50[2)] |
| B1 | 1036.6717 | 560.36 | 478.33 | 236.17[2)] |
| **A2*** | **1036.6604** | **280.67**[1)] | **478.33** | **231.50**[2)] |
| **A1*** | **1050.6761** | **574.33** | **478.33** | **236.17**[2)] |
| **B2*** | **1064.6666** | **588.33** | **478.33** | **231.50**[2)] |
| **B1*** | **1078.6822** | **602.36** | **478.33** | **236.17**[2)] |

[1)] ($z$=2), [2)] ($z$=3)

**Figure S18. $MS^2$ fragmentation of different paenilamicin and *N*-acetylpaenilamicin variants to determine regioselective acetylation.** Observed fragment ions $b_4$, $y_4$ and $y_6$ are highlighted in the chemical structure of paenilamicin. The different paenilamicin variants with residue $R_1$ (Glm, Aga) and $R_2$ (Orn, Lys) are shown as circles. Fragment ions of *N*-acetylpaenilamicin variants are indicated bold (magenta) and with an asterisk. The schematic $MS^2$ spectrum shows the observed fragment ions of each single paenilamicin and *N*-acetylpaenilamicin variant. Arrows show distinct mass shifts of acetylation (magenta) and the different paenilamicin series (blue and green). Observed fragment ions are listed as mass-to-charge ratios with $z$=1.

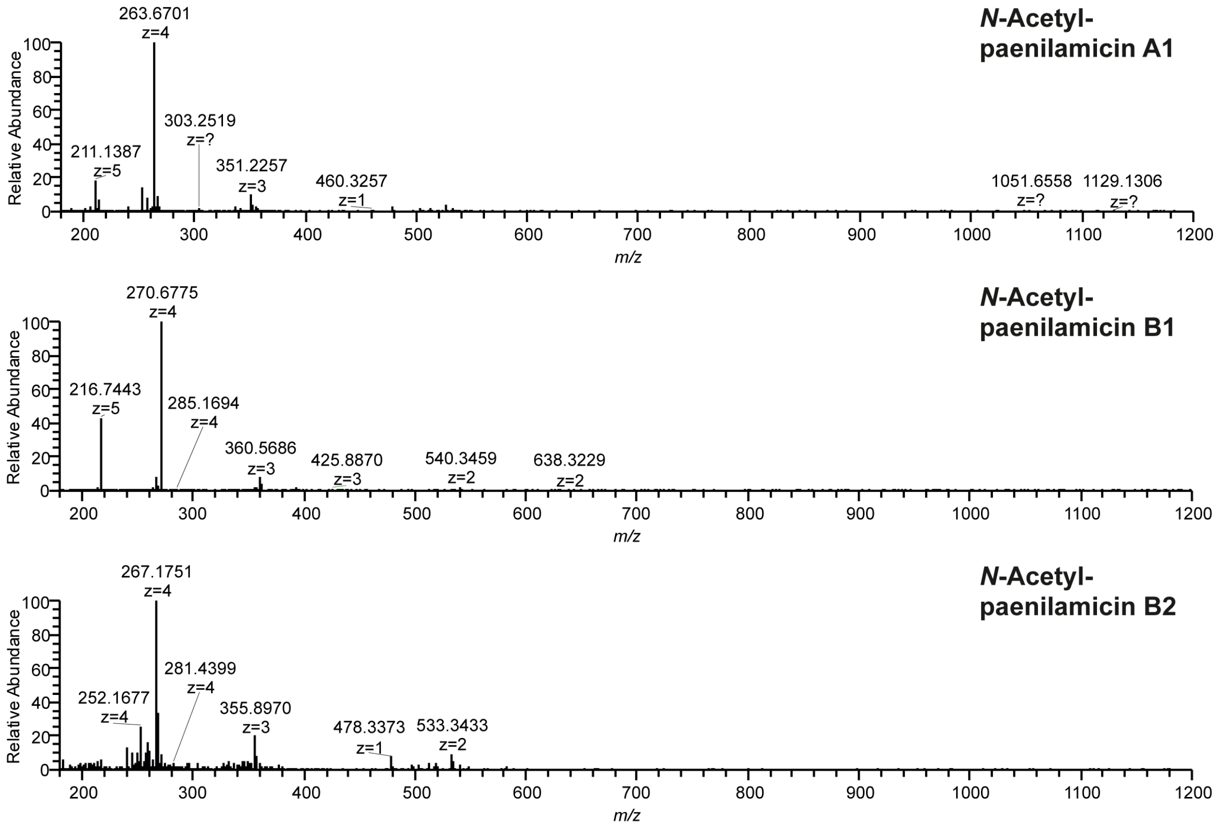


**Figure S19.** $MS^1$ spectra of *N*-acetylpaenilamicin A1, B1 and B2 isolated from *P. larvae*.

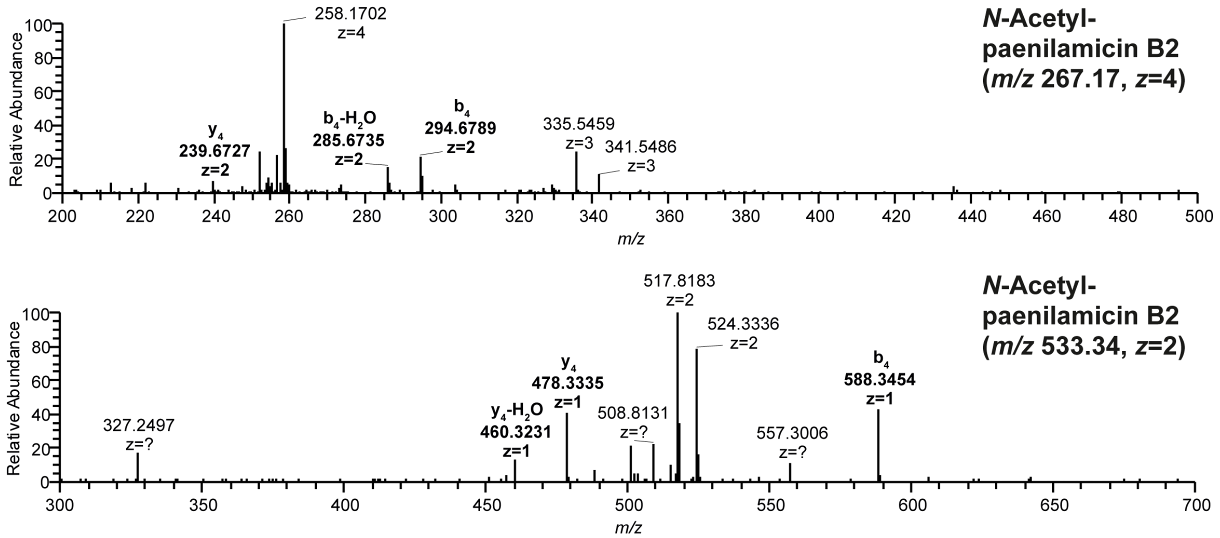


**Figure S20.** $MS^2$ spectra of *N*-acetylpaenilamicin B2 isolated from *P. larvae* DSM 25430 (ERIC II).

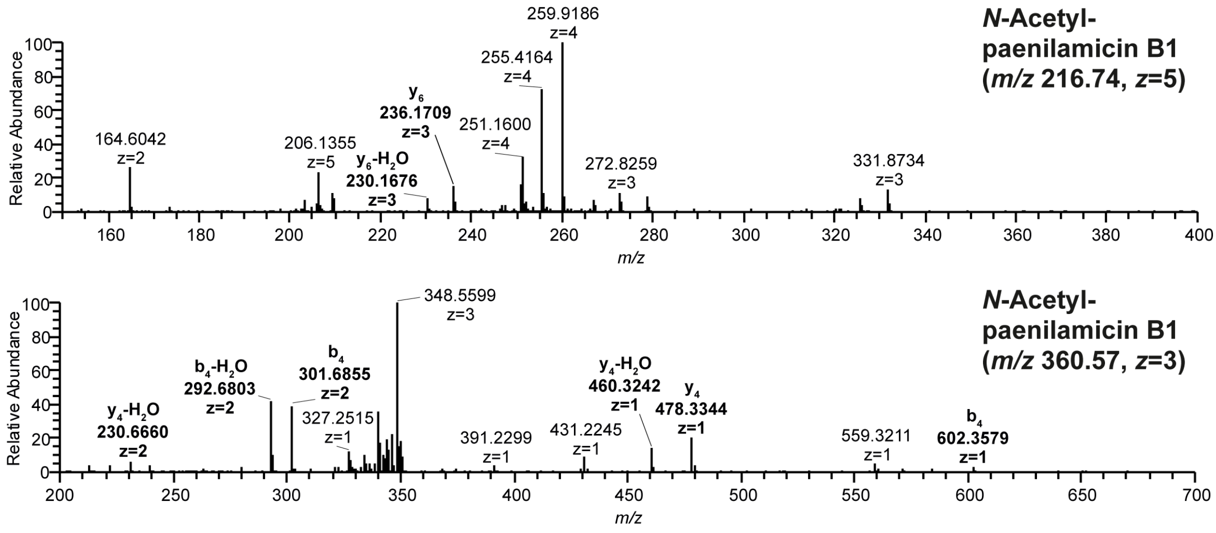


**Figure S21.** $MS^2$ spectra of *N*-acetylpaenilamicin B1 isolated from *P. larvae* DSM 25430 (ERIC II).

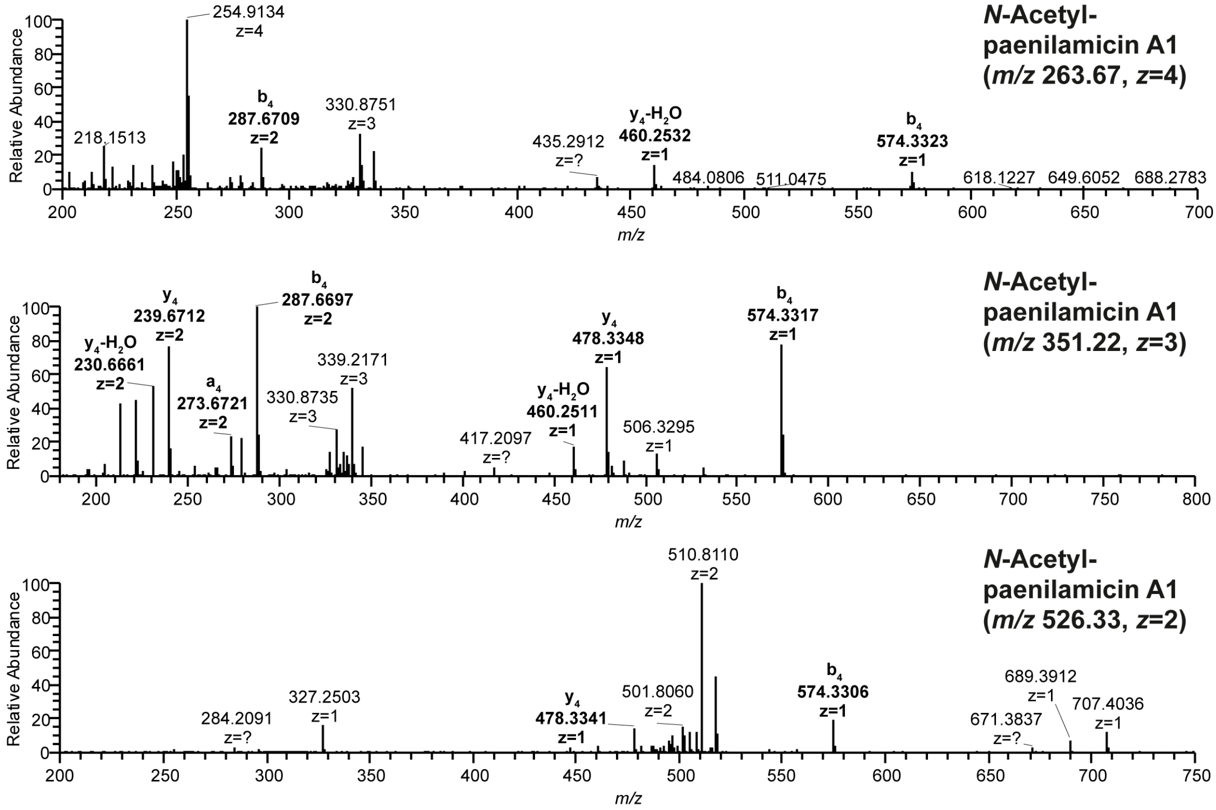


**Figure S22.** $MS^2$ spectra of *N*-acetylpaenilamicin A1 isolated from *P. larvae* ATCC 9545 (ERIC I).

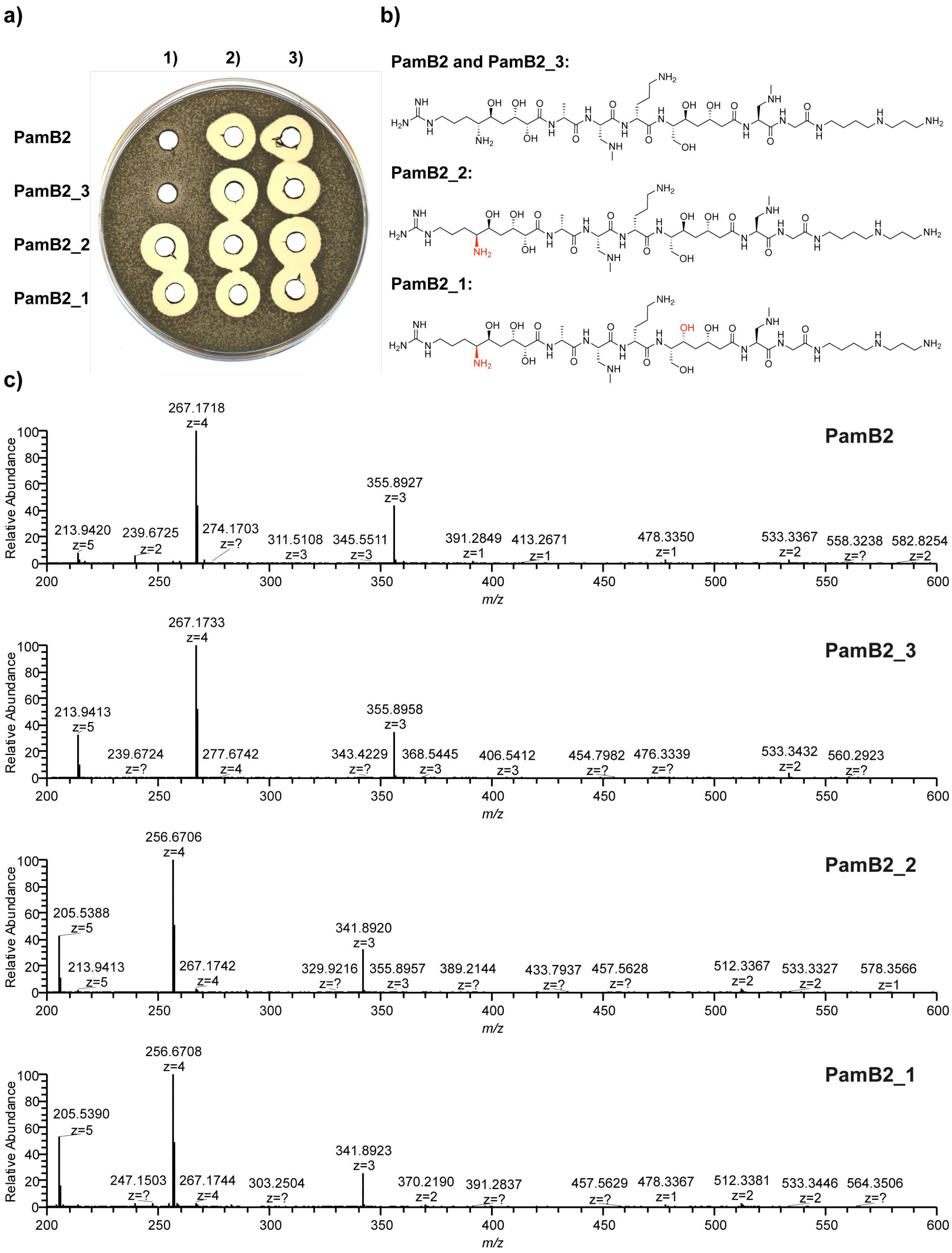


**Figure S23. Stereoselective *N*-acetylation of PamZ.** a) Natural paenilamicin B2 (PamB2), synthetic paenilamicin B2 (PamB2_3) and its diastereomers (PamB2_2, PamB2_1) were incubated with acetyl-CoA and PamZ (1), only acetyl-CoA (2), only PamZ (3) *in vitro* and tested by agar diffusion assay against *B. megaterium* as indicator strain. b) Chemical structure of paenilamicin B2 and its diastereomers. c) *N*-acetylation observed and verified by HPLC-ESI MS.

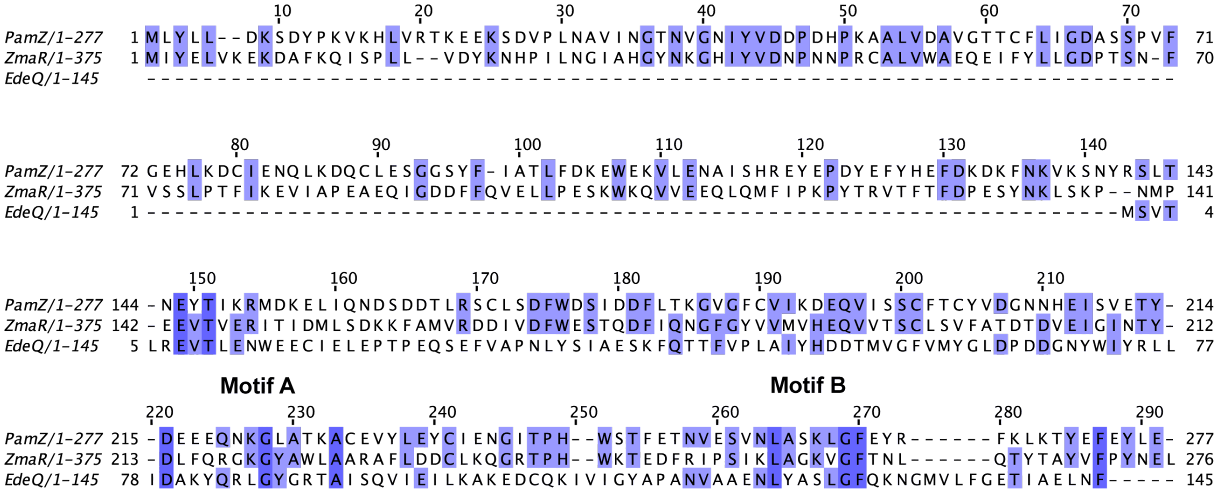


**Figure S24. Multiple sequence alignment of Gcn5-related *N*-acetyltransferases of PamZ (paenilamicin), ZmaR (zwittermicin) and EdeQ (edeine).** Blue shaded residues are identical. The pyrophosphate group of coenzyme A interacts with the highly conserved motif A and the adenine ring with the not well conserved motif B.

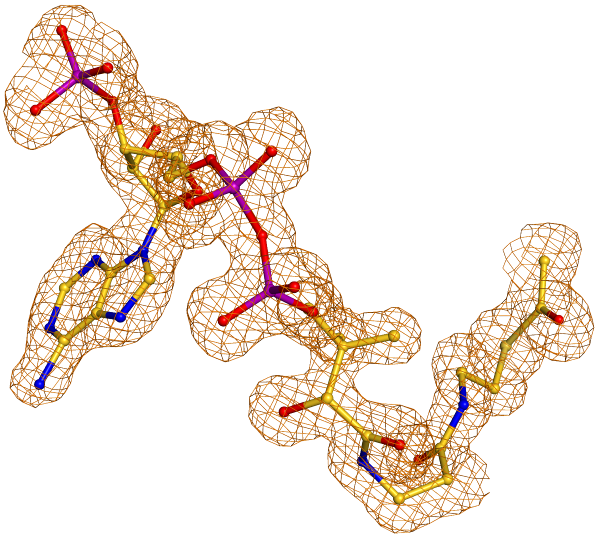

**Figure S25.** Polder electron density map[1] of acetyl-CoA shown as mesh at a σ-level of 3.0. Acetyl-CoA is presented as ball-stick-model with carbon atoms colored in magenta, oxygen in red, phosphorous in orange and nitrogen in light blue. Hydrogen atoms are not shown.

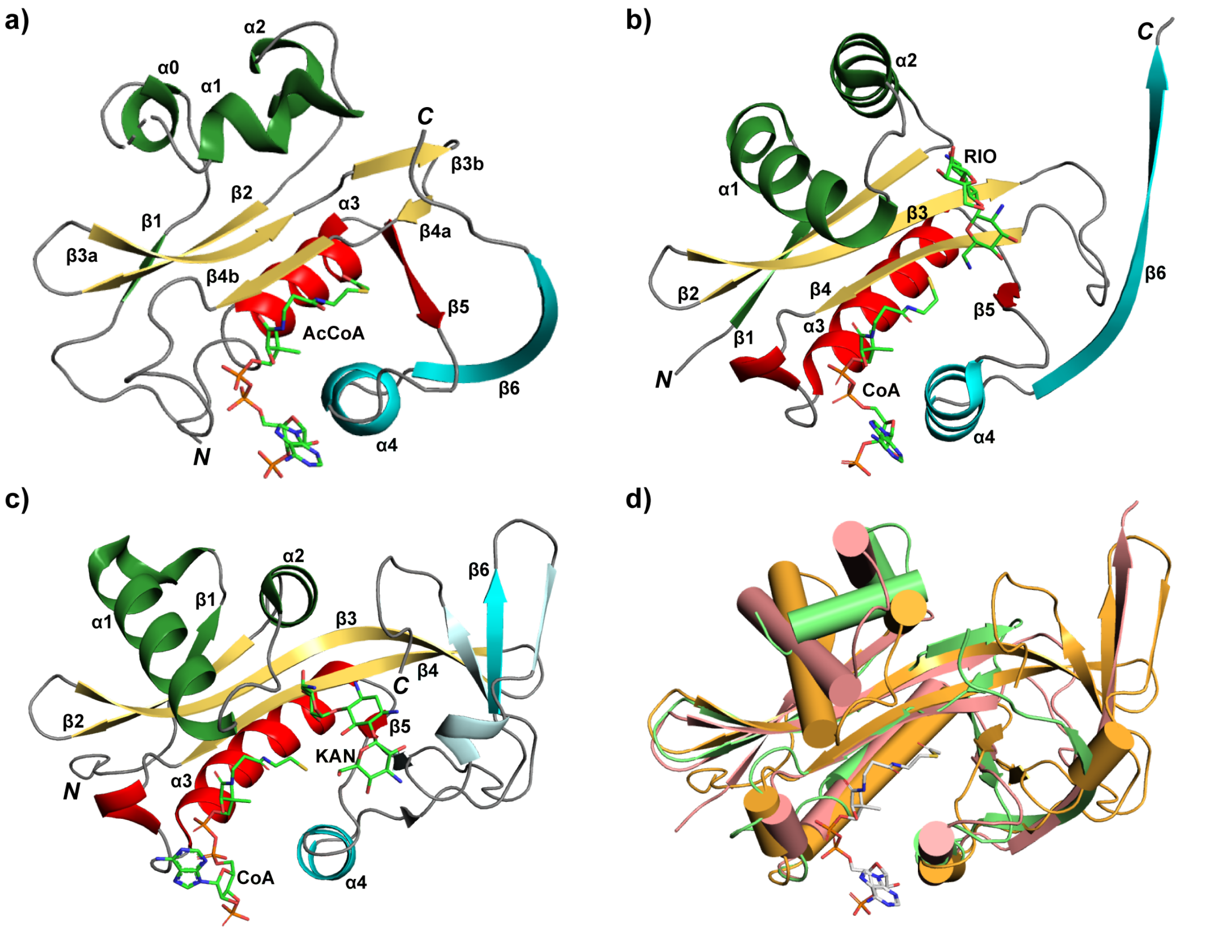


**Figure S26. Structural comparison of PamZ and AACs.** a) C-terminal domain of PamZ including acetyl-CoA, b) AAC(6') from *Salmonella enterica* including CoA and ribostamycin (PDB ID: 1s3z), c) AAC(2') from *Mycobacterium tuberculosis* including CoA and kanamycin A (PDB ID: 1m4i), d) pair-wise structural alignment of C-terminal domain of PamZ (green), AAC(6') (pink) and AAC(2') (orange). The CTD of PamZ superimposes with the AAC(6') with an RMSD of 1.9 Å for 104 pairs of Cα-atoms. AAC(2') superimposes with CTD of PamZ with an RMSD of 4.0 Å for 75 pairs of Cα-atoms.

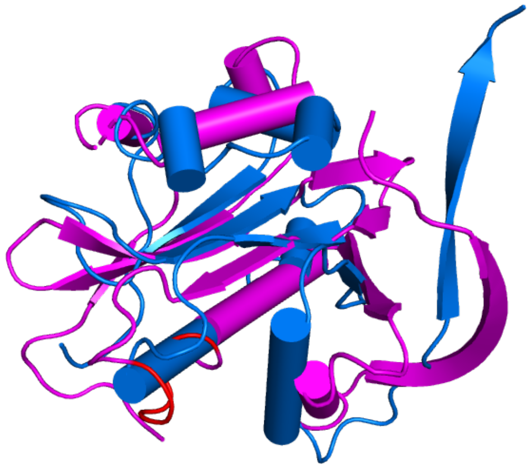

**Figure S27. Pairwise structural alignment between NTD and CTD of PamZ.** The NTD is highlighted in magenta and CTD in blue. The P-loop only present in CTD is highlighted in red.

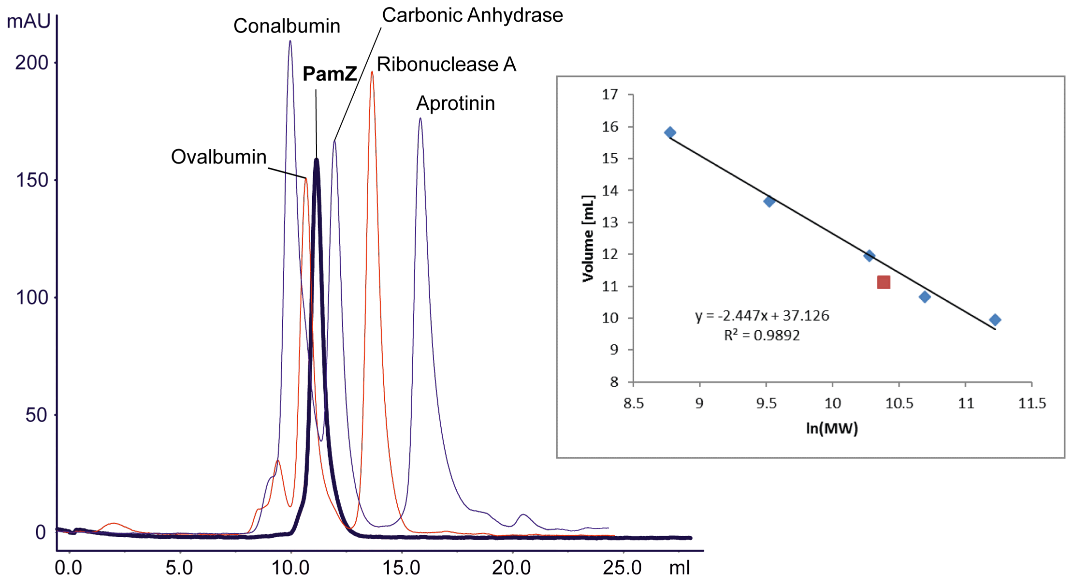


**Figure S28. Size exclusion chromatogram and calibration curve of PamZ including mixture A and B as calibration standards detected at 280 nm.** Chromatogram indicates PamZ as monomeric unit. The calibration standards (blue squares) have the following molecular weights: conalbumin (75 kDa), ovalbumin (44 kDa), carbonic anhydrase (29 kDa), ribonuclease A (13.7 kDa), aprotinin (6.5 kDa).

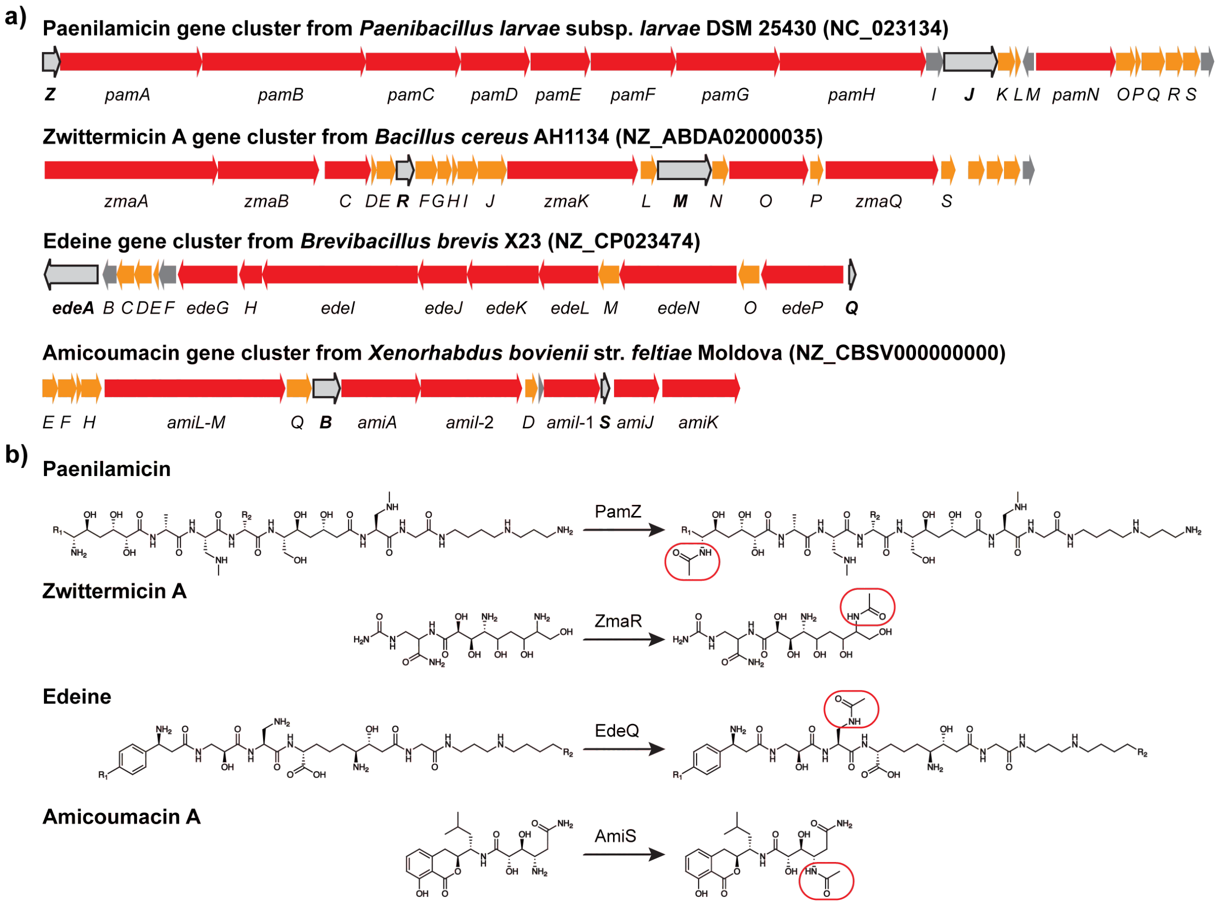


**Figure S29. Biosynthetic gene cluster and *N*-acetylation of paenilamicin, zwittermicin A, edeine and amicoumacin.** a) Different functions of the genes are grouped by colour: core biosynthetic gene (red), additional biosynthetic gene (orange), other gene (dark grey), resistance gene (light grey and framed in black). b) *N*-acetylation reaction of the cationic peptides. Position of *N*-acetylation framed in red.

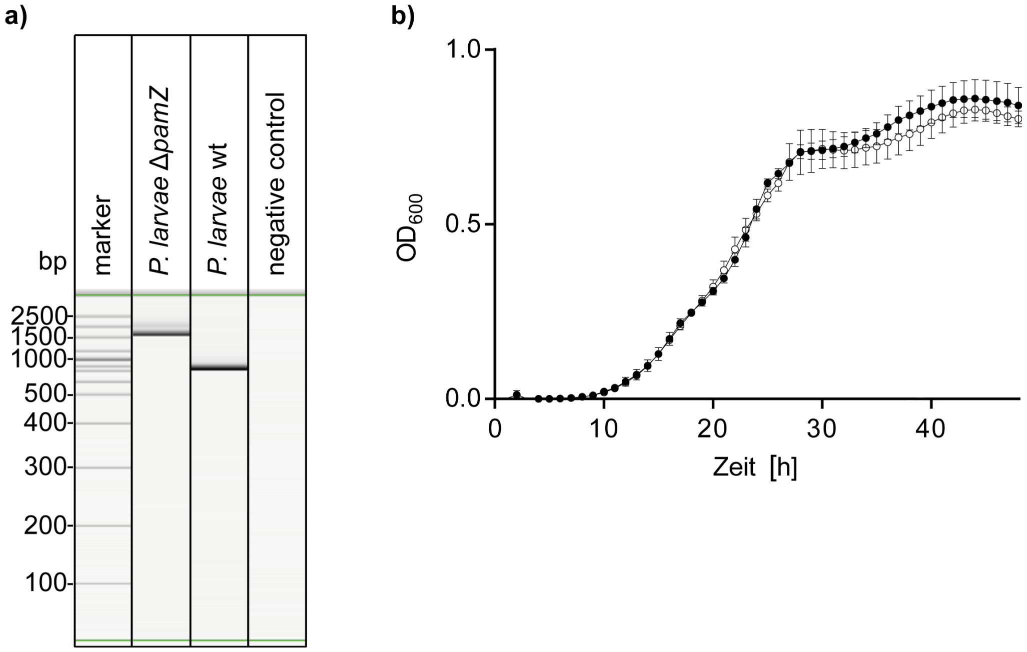


**Figure S30. Determination of successful intron insertion into the *pamZ* gene of *P. larvae* DSM 25430.** a) PCR analysis with gel electrophoresis revealed an about 900 bp larger fragment caused by intron insertion in the knockout mutant Δ*pamZ* (1612 bp) compared with the wild type strain (712 bp). b) Growth curves of *P. larvae* WT (black circles) and the *P. larvae* Δ*pamZ* deletion mutant (white circles) in MYPGP liquid broth under anaerobic conditions (two-way-ANOVA, p=0.6486).

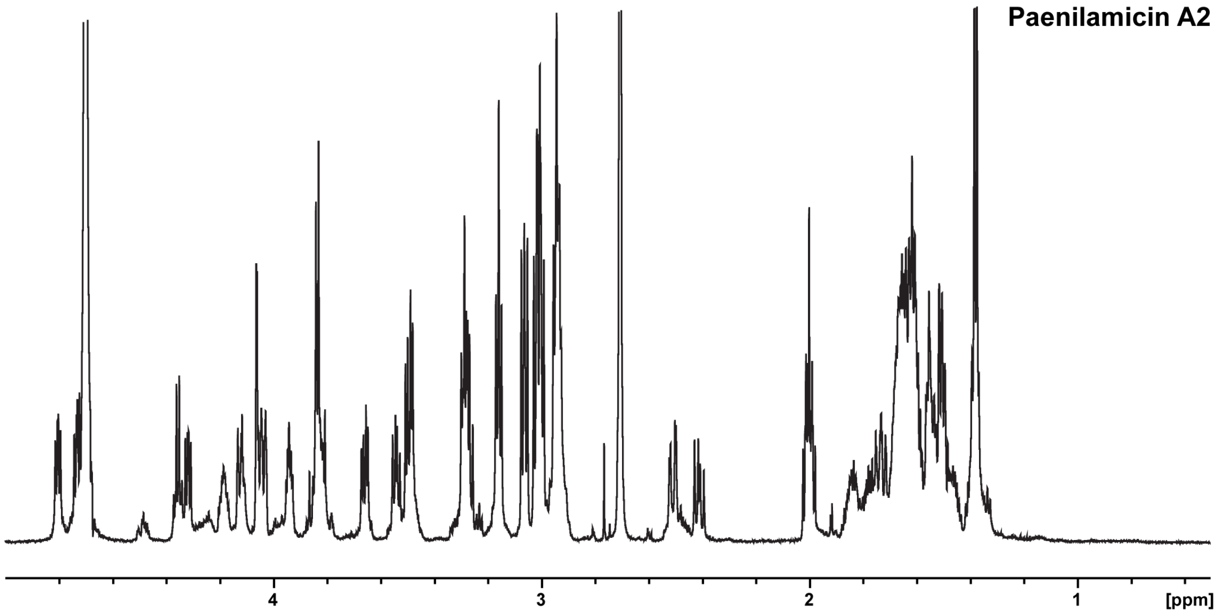


**Figure S31.** $^1$H-NMR spectrum of paenilamicin A2 isolated from *P. larvae* ATCC 9545 and recorded in $D_2O$ at 298 K displays aliphatic region.

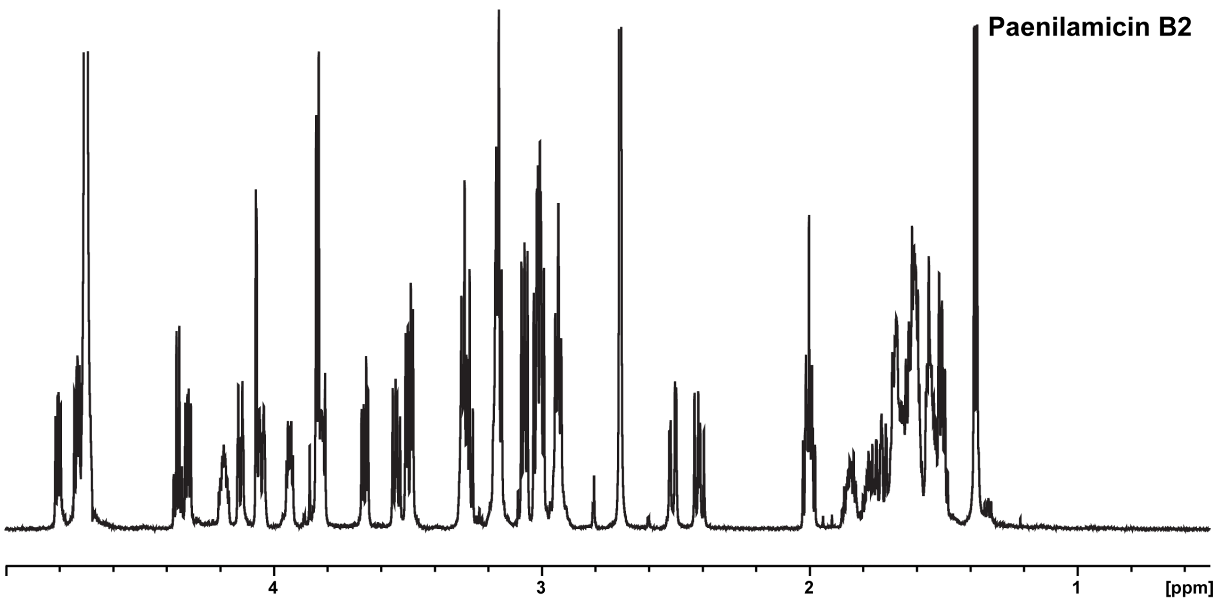


**Figure S32.** [1]H-NMR spectrum of paenilamicin B2 isolated from *P. larvae* ATCC 9545 and recorded in $D_2O$ at 298 K displays aliphatic region.

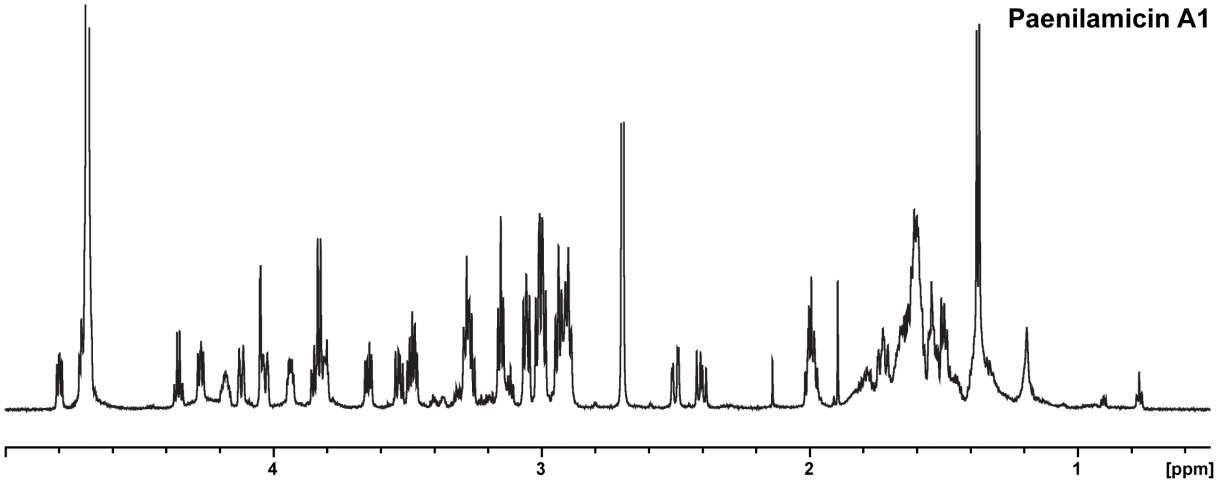


**Figure S33.** [1]H-NMR spectrum of paenilamicin A1 isolated from *P. larvae* DSM 25430 and recorded in $D_2O$ at 298 K displays aliphatic region.

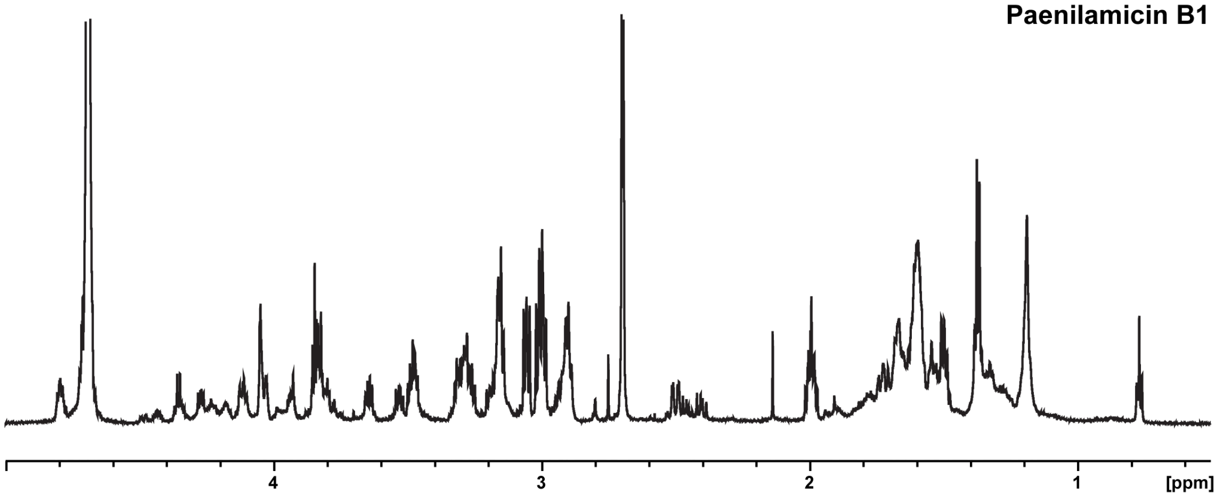


**Figure S34.** $^{1}$H-NMR spectrum of paenilamicin B1 isolated from *P. larvae* DSM 25430 and recorded in $D_2O$ at 298 K displays aliphatic region.

### Supplementary Tables

**Table S1. Fragment ions of paenilamicin, *N*-acetylpaenilamicin and *N*²-acetylpaenilamicin variants.**

**Table S2. Comparison of $^{1}$H and $^{13}$C chemical shifts of paenilamicin B2 and *N*-acetylpaenilamicin B2 in $D_2O$ and 0.1% acetic acid-$d_4$ at 298 K.**

**Table S3. Crystallographic data collection and model refinement statistics.**

**Table S4. Primers used for knockout generation and screening for knockout mutants.**

**Table S5. Oligonucleotides used in this study.**

**Table S1. Fragment ions of paenilamicin, *N*-acetylpaenilamicin and *N*²-acetylpaenilamicin variants.**

| Compound | Observed *m/z* | Calculated *m/z* | Fragment ions |
|---|---|---|---|
| Paenilamicin A1 (isolated) | *m/z* 337.23 (*z*=3)<br>*m/z* 505.34 (*z*=2) | *m/z* 337.23 (*z*=3)<br>*m/z* 505.34 (*z*=2) | $b_4$, $y_4$<br>$b_4$, $y_4$, $y_6$, $y_7$ |
| Paenilamicin A2 (isolated) | *m/z* 332.56 (*z*=3)<br>*m/z* 498.33 (*z*=2) | *m/z* 332.56 (*z*=3)<br>*m/z* 498.33 (*z*=2) | $b_4$, $y_4$<br>$b_4$, $y_4$, $y_6$, $y_7$ |
| Paenilamicin B1 (isolated) | *m/z* 208.34 (*z*=5)<br>*m/z* 346.56 (*z*=3) | *m/z* 208.34 (*z*=5)<br>*m/z* 346.56 (*z*=3) | $b_4$, $y_4$, $y_6$<br>$b_4$, $y_4$ |
| Paenilamicin B2 (isolated) | *m/z* 205.54 (*z*=5)<br>*m/z* 256.67 (*z*=4)<br>*m/z* 341.89 (*z*=3)<br>*m/z* 512.33 (*z*=2) | *m/z* 205.54 (*z*=5)<br>*m/z* 256.67 (*z*=4)<br>*m/z* 341.89 (*z*=3)<br>*m/z* 512.34 (*z*=2) | $b_4$, $y_4$, $y_6$<br>$b_4$, $y_2$, $y_4$<br>$b_4$, $y_4$<br>$b_4$, $y_4$ |
| Paenilamicin B2 (synthesized) | *m/z* 205.54 (*z*=5)<br>*m/z* 341.89 (*z*=3) | *m/z* 205.54 (*z*=5)<br>*m/z* 341.89 (*z*=3) | $b_4$, $y_4$, $y_6$<br>$b_4$, $y_4$ |
| *N*-acetylpaenilamicin A1 (isolated, *in vitro*) | *m/z* 211.14 (*z*=5) | *m/z* 211.14 (*z*=5) | $y_6$ |
| *N*²-acetylpaenilamicin A1 (isolated, *in vitro*) | *m/z* 365.24 (*z*=3) | *m/z* 365.24 (*z*=3) | $b_4$, $y_4$ |
| *N*-acetylpaenilamicin A2 (isolated, *in vitro*) | *m/z* 208.34 (*z*=5)<br>*m/z* 346.56 (*z*=3) | *m/z* 208.34 (*z*=5)<br>*m/z* 346.56 (*z*=3) | $b_4$, $y_4$, $y_6$<br>$b_4$, $y_4$ |
| *N*²-acetylpaenilamicin A2 (isolated, *in vitro*) | *m/z* 360.57 (*z*=3) | *m/z* 360.56 (*z*=3) | $b_4$, $y_4$ |
| *N*-acetylpaenilamicin B1 (isolated, *in vitro*) | *m/z* 216.74 (*z*=5)<br>*m/z* 360.57 (*z*=3) | *m/z* 216.74 (*z*=5)<br>*m/z* 360.57 (*z*=3) | $y_6$<br>$b_4$, $y_4$ |
| *N*-acetylpaenilamicin B2 (isolated, *in vitro*) | *m/z* 213.94 (*z*=5)<br>*m/z* 267.17 (*z*=4)<br>*m/z* 355.89 (*z*=3) | *m/z* 213.94 (*z*=5)<br>*m/z* 267.17 (*z*=4)<br>*m/z* 355.90 (*z*=3) | $b_4$, $y_4$, $y_6$<br>$b_4$<br>$a_4$, $b_4$, $y_4$ |
| *N*-acetylpaenilamicin B2 (synthesized, *in vitro*) | *m/z* 213.94 (*z*=5)<br>*m/z* 267.17 (*z*=4) | *m/z* 213.94 (*z*=5)<br>*m/z* 267.17 (*z*=4) | $b_4$, $y_4$, $y_6$<br>$b_4$ |
| *N*-acetylpaenilamicin A1 (isolated) | *m/z* 263.67 (*z*=4)<br>*m/z* 351.22 (*z*=3)<br>*m/z* 526.33 (*z*=2) | *m/z* 263.68 (*z*=4)<br>*m/z* 351.23 (*z*=3)<br>*m/z* 526.35 (*z*=2) | $b_4$, $y_4$<br>$a_4$, $b_4$, $y_4$<br>$b_4$, $y_4$ |
| *N*-acetylpaenilamicin B1 (isolated) | *m/z* 216.74 (*z*=5)<br>*m/z* 360.57 (*z*=3) | *m/z* 216.74 (*z*=5)<br>*m/z* 360.57 (*z*=3) | $y_6$<br>$b_4$, $y_4$ |

| | | | |
|---|---|---|---|
| *N*-acetylpaenilamicin B2 (isolated) | *m/z* 267.17 (*z*=4)<br>*m/z* 533.34 (*z*=2) | *m/z* 267.17 (*z*=4)<br>*m/z* 533.34 (*z*=2) | $b_4$, $y_4$<br>$b_4$, $y_4$ |

**Table S2. Comparison of $^1$H and $^{13}$C chemical shifts of paenilamicin B2 and *N*-acetylpaenilamicin B2 in $D_2O$ and 0.1% acetic acid-$d_4$ at 298 K.** Chemical shifts and chemical shift perturbations (CSPs) are given in ppm. The color code refers to that used in Figure 3b of the main manuscript.

| | | Paenilamicin B2 | | *N*-acetylpaenilamicin B2 | | |
|---|---|---|---|---|---|---|
| **Residue** | **Pos.** | **δ $^1$H** | **δ $^{13}$C** | **δ $^1$H** | **δ $^{13}$C** | **CSP** |
| **Aga** | **2** | 4.15 | 77.26 | 4.13 | 77.40 | 0.03 |
| | **3** | 4.21 | 71.00 | 4.20 | 71.35 | 0.03 |
| | **4'** | 1.82 | 37.78 | 1.82 | 38.66 | 0.05 |
| | **4''** | 1.63 | | 1.52 | | 0.12 |
| | **5** | 4.13 | 69.78 | 3.77 | 72.41 | 0.39 |
| | **6** | 3.38 | 58.64 | 3.82 | 56.76 | 0.45 |
| | **7'** | 1.74 | 26.76 | 1.74 | 28.82 | 0.12 |
| | **7''** | 1.67 | | 1.44 | | 0.26 |
| | **8'** | 1.75 | 27.22 | 1.66 | 27.34 | 0.09 |
| | **8''** | 1.67 | | 1.56 | | 0.10 |
| | **9** | 3.26 | 43.38 | 3.20 | 43.49 | 0.06 |
| **Ac** | **2** | n.d.[a] | n.d.[a] | 2.04 | 24.71 | n.d.[a] |
| **Ala** | α | 4.44 | 52.38 | n.d.[a] | n.d.[a] | n.d.[a] |
| | β | 1.47 | 19.06 | 1.46 | 18.90 | 0.01 |
| **mDap1** | α | 4.82 | 52.73 | n.d.[a] | n.d.[a] | n.d.[a] |
| | β' | 3.58 | 51.63 | 3.58 | 51.63 | 0.00 |
| | β'' | 3.38 | | 3.38 | | 0.00 |
| | δ | 2.79 | 36.20 | 2.79 | 36.18 | 0.00 |
| **Orn** | α | 4.41 | 56.52 | n.d.[a] | n.d.[a] | n.d.[a] |
| | β' | 1.94 | 30.86 | 1.94 | 30.85 | 0.00 |
| | β'' | 1.87 | | 1.87 | | 0.00 |
| | γ | 1.74 | 26.04 | 1.73 | 26.02 | 0.01 |
| | δ | 3.03 | 41.58 | 3.02 | 41.59 | 0.00 |
| **Gla** | **2'** | 2.59 | 46.36 | 2.60 | 46.35 | 0.00 |
| | **2''** | 2.51 | | 2.50 | | 0.01 |
| | **3** | 4.27 | 68.03 | 4.27 | 68.02 | 0.00 |
| | **4** | 1.64 | 43.05 | 1.64 | 43.08 | 0.00 |
| | **5** | 4.03 | 68.97 | 4.03 | 68.95 | 0.01 |
| | **6** | 3.91 | 58.77 | 3.90 | 58.77 | 0.01 |
| | **7'** | 3.75 | 63.76 | 3.74 | 63.79 | 0.01 |
| | **7''** | 3.63 | | 3.63 | | 0.00 |
| **mDap2** | α | 4.89 | 52.43 | n.d.[a] | n.d.[a] | n.d.[a] |
| | β' | 3.58 | 51.81 | 3.58 | 51.81 | 0.00 |
| | β'' | 3.36 | | 3.36 | | 0.00 |
| | δ | 2.79 | 36.20 | 2.79 | 36.18 | 0.00 |
| **Gly** | α | 3.92 | 45.24 | 3.92 | 45.22 | 0.00 |
| **Spd** | **2** | 3.11 | 39.27 | 3.10 | 39.27 | 0.00 |
| | **3** | 2.09 | 26.45 | 2.09 | 26.48 | 0.01 |
| | **4** | 3.15 | 47.18 | 3.15 | 47.19 | 0.00 |
| | **6** | 3.09 | 50.07 | 3.09 | 50.07 | 0.00 |

| | | | | | |
|---|---|---|---|---|---|
| **7** | 1.70 | 25.69 | 1.71 | 25.69 | 0.00 |
| **8** | 1.59 | 28.20 | 1.59 | 28.24 | 0.00 |
| **9** | 3.25 | 41.36 | 3.25 | 41.37 | 0.00 |

[a] not determined.

**Table S3. Crystallographic data collection and model refinement statistics for binary complex PamZ incl. acetyl-CoA.**

| **Data Collection** | |
|---|---|
| PDB ID | 7B3A |
| Wavelength [Å] | 0.91841 |
| Temperature [K] | 100 |
| Space group | $P2_1$ |
| **Unit cell parameters** | |
| a, b, c [Å] | 36.4, 70.1, 54.6 |
| $\alpha$, $\beta$, $\gamma$ [°] | 90.0, 107.0, 90.0 |
| Resolution range [Å][a] | 50.00-1.34 (1.42-1.34) |
| **Reflections[a]** | |
| Unique | 58,898 (9,129) |
| Completeness [%] | 98.2 (94.5) |
| Multiplicits | 3.8 (3.7) |
| **Data quality[a]** | |
| Intensity [I/σ(I)] | 11.20 (1.00) |
| $R_{meas}$ [%][b] | 6.8 (129.2) |
| $CC_{1/2}$[c] | 99.9 (37.9) |
| Wilson B value [$Å^2$] | 23.1 |
| **Refinement** | |
| Resolution range [Å ][a] | 50.00-1.34 (1.37-1.34) |
| **Reflections[a]** | |
| Number | 58,882 (3,578) |
| Test set (3.6%) | 2,099 (127) |
| $R_{work}$[a] | 0.147 (0.305) |
| $R_{free}$[a] | 0.181 (0.358) |
| **Contents of the asymmetric unit** | |
| Protein, molecules, residues, atoms | 1, 1, 431, 3774 |
| Acetyl-CoA, acetate molecules, chloride | 1, 1, 1 |
| Water, molecules | 270 |
| **Mean temperature factors [$Å^2$][b]** | |
| All Atoms | 24.5 |
| Macromolecules | 23.8 |
| Ligands | 28.0 |
| Water oxygens | 32.5 |
| **RMSD[d] from target geometry** | |
| Bond lengths [Å] | 0.016 |
| Bond angles [°] | 1.500 |
| **Validation statistics[c]** | |
| Ramachandran plot[e] | |

| | |
|---|---|
| Residues in allowed regions [%] | 1.8 |
| Residues in favored regions [%] | 98.2 |
| Rotamer outliers [%][e] | 1.0 |
| MOLPROBITY clashscore[f,g] | 3.0 |
| MOLPROBITY overall[f] | 1.1 |

[a] data for the highest resolution shell in parenthesis.

[b] $R_{meas}(I) = \sum_h [N/(N-1)]^{1/2} \sum_i \left| I_{ih} - \langle I_h \rangle \right| / \sum_h \sum_i I_{ih}$, in which $\langle I_h \rangle$ is the mean intensity of symmetry-equivalent reflections h, $I_{ih}$ is the intensity of a particular observation of h and N is the number of redundant observations of reflection h.[2]

[c] $CC_{1/2} = (\langle I^2 \rangle - \langle I \rangle^2) / (\langle I^2 \rangle - \langle I \rangle^2) + \sigma^2_\varepsilon$, in which $\sigma^2_\varepsilon$ is the mean error within a half-dataset.[3]

[d] Root mean square deviation.

[e] calculated with PHENIX.[4]

[f] calculated with MOLPROBITY.[5]

[g] clashscore is the number of serious steric overlaps (> 0.4) per 1,000 atoms.[5]

**Table S4. Primers used for knockout generation and screening for knockout mutants.**

| Name | Sequence 5'→3' | Site |
|---|---|---|
| pamZ_118_IBS | AAAAAAGCTTATAATTATCCTTACATCTCCATAAAGTGCGCCCAGA-TAGGGTG | Knockout generation |
| pamZ_118_EBS1 | CAGATTGTACAAATGTGGTGATAACAGATAAGTCCATAAA-TATAACTTACCTTTCTTTGT | Knockout generation |
| pamZ_118_EBS2 | TGAACGCAAGTTTCTAATTTCGGTTAGATGTCGATAGAG-GAAAGTGTCT | Knockout generation |
| pamZ_fw | GTGATGTGCCTCTCAATGCAG | Knockout screening |
| pamZ_rev | CCCAGCTTACTTGCCAGGTT | Knockout screening |

**Table S5. Oligonucleotides used in this study.**

| Name | Sequence 5'→3' | Site |
|---|---|---|
| pET28a_pamZ_for | GGCCATATGGCsAGCATGCTGTATCTATTGGATAAATCGG | *Nhe*I |
| pET28a_pamZ_rev | CTCGAGTCTCGAGTTAA-TATTCAAATTCATAGGTTTTAAGTTTAAACC | *Xho*I |
